# Simulations support the interaction of the SARS-CoV-2 spike protein with nicotinic acetylcholine receptors

**DOI:** 10.1101/2020.07.16.206680

**Authors:** A. Sofia F. Oliveira, Amaurys Avila Ibarra, Isabel Bermudez, Lorenzo Casalino, Zied Gaieb, Deborah K. Shoemark, Timothy Gallagher, Richard B. Sessions, Rommie E. Amaro, Adrian J. Mulholland

## Abstract

Changeux *et al*. recently suggested that the SARS-CoV-2 spike (S) protein may interact with nicotinic acetylcholine receptors (nAChRs). Such interactions may be involved in pathology and infectivity. Here, we use molecular simulations of validated atomically detailed structures of nAChRs, and of the S protein, to investigate this ‘nicotinic hypothesis’. We examine the binding of the Y674-R685 loop of the S protein to three nAChRs, namely the human α4β2 and α7 subtypes and the muscle-like αβγδ receptor from *Tetronarce californica*. Our results indicate that Y674-R685 has affinity for nAChRs and the region responsible for binding contains the PRRA motif, a four-residue insertion not found in other SARS-like coronaviruses. In particular, R682 has a key role in the stabilisation of the complexes as it forms interactions with loops A, B and C in the receptor’s binding pocket. The conformational behaviour of the bound Y674-R685 region is highly dependent on the receptor subtype, adopting extended conformations in the α4β2 and α7 complexes and more compact ones when bound to the muscle-like receptor. In the α4β2 and αβγδ complexes, the interaction of Y674-R685 with the receptors forces the loop C region to adopt an open conformation similar to other known nAChR antagonists. In contrast, in the α7 complex, Y674-R685 penetrates deeply into the binding pocket where it forms interactions with the residues lining the aromatic box, namely with TrpB, TyrC1 and TyrC2. Estimates of binding energy suggest that Y674-R685 forms stable complexes with all three nAChR subtypes. Analyses of the simulations of the full-length S protein show that the Y674-R685 region is accessible for binding, and suggest a potential binding orientation of the S protein with nAChRs.

## Main text

The severe acute respiratory syndrome coronavirus-2 (SARS-CoV-2) is a novel strain of coronavirus that first appeared in China in late 2019 and causes the potentially fatal disease COVID-19. This virus initially infects respiratory epithelial cells by binding to the angiotensin-converting 2 enzyme^1^ (ACE2) and may cause pneumonia and severe acute respiratory distress syndrome.^2,3^ Since it emerged as a human pathogen, SARS-CoV-2 has caused more than 27.5 million confirmed cases of COVID-19 and 903,756 deaths worldwide as of 9^th^ September, 2020.^4^ Several major risk factors for the development of COVID-19 have been identified, namely age, heart disease, diabetes and hypertension.^5^ Recently, given the apparently low prevalence of smokers among hospitalised COVID-19 patients,^6-8^ it was proposed that nicotine may offer some protective value to mitigate COVID-19 (the ‘protection’ hypothesis).^6^ It has been suggested that medicinal nicotine (either in patches, gum, or electronic delivery systems) should be investigated as a therapeutic option for this disease.^6,9^ Clinical trials for nicotine are underway (e.g. https://clinicaltrials.gov/ct2/show/NCT04429815). It should be noted that alternative explanations to the ‘protection’ hypothesis have been proposed:^10^ the first relates to the failure in correctly identifying smokers upon hospital admission,^10^ and the second is that hospitalised COVID-19 patients may be less likely to smoke as their comorbidities motivate them to quit (‘smoking cessation’ hypothesis).^10^

Based on the early observations of the lower than expected smoking prevalence in hospitalised COVID-19 patients, Changeux and colleagues suggested a role for nicotinic acetylcholine receptors (nAChRs) in the pathophysiology of COVID-19 via a direct interaction between these receptors and the viral spike (S) glycoprotein.^11^ This suggestion was based in the fact that the S protein from SARS-CoV-2 contains a sequence motif similar to known nAChR antagonists^11^ (**Figure S1**), such as α-bungarotoxin from *Bungarus multicinctus* and glycoprotein from *Rabies lyssavirus* (formerly *Rabies virus*). Changeux *et al*. also proposed that COVID-19 might be controlled or mitigated by the use of nicotine, if the latter can sterically or allosterically compete with the virus for binding to these receptors.^9,11^

nAChRs are cation channels that belong to the pentameric ligand-gated ion channel family.^12^ They are present in both the peripheral (at the skeletal neuromuscular junction and in the autonomic nervous system) and central nervous system (CNS).^13^ The neuronal receptors have emerged as important targets for the treatment of Alzheimer’s disease, schizophrenia, pain and nicotine addiction.^13,14^ Mutations of muscle nAChR can cause congenital myasthenia gravis.^4^ There is a large repertoire of nAChR subtypes which differ in the homo-or heteromeric assembly of five monomers arranged around a central channel axis.^15-17^ Each subtype shows different selectivity for agonists and antagonists.^15-17^ All nAChRs share the same basic architecture (**Figure 1B**), formed of a large N-terminal extracellular domain (ECD), where the agonist binding site is located; a transmembrane domain (TMD) surrounding the ion channel; an intracellular domain (ICD); and a short extracellular C-terminal domain (CTD).^15-17^ The ligand-binding pocket is located at the interface between two neighbouring subunits (**Figure 1B**) and is formed by loops A, B and C from the principal subunit and D, E and F from the complementary subunit (**Figure S2**).

**Figure 1.**
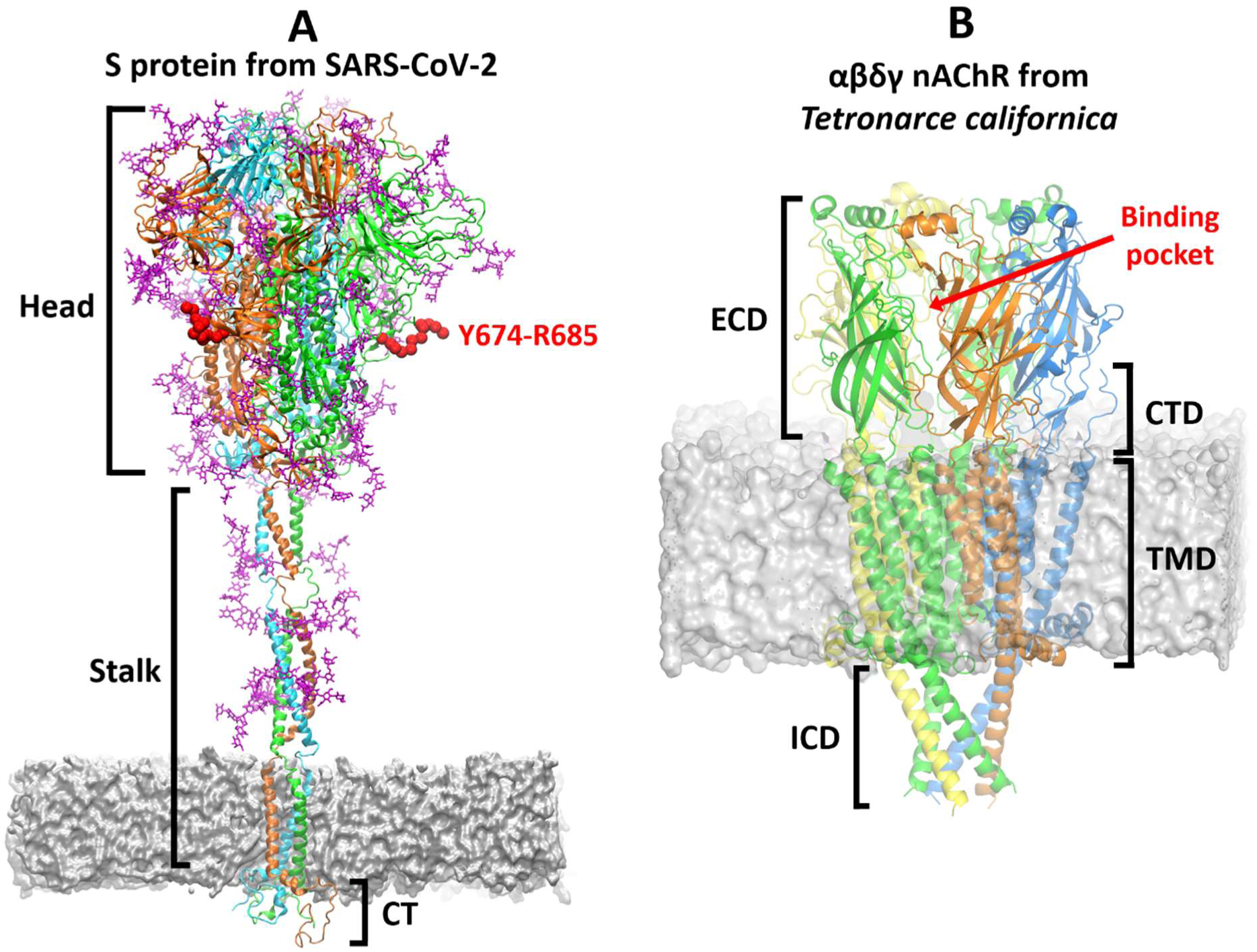
Overview of the three-dimensional structures of the S protein from SARS-CoV-2 and the αβγd nAChR from *Tetronarce californica*. (**A)** The model for the complete, fully glycosylated, SARS-CoV-2 S protein represents the closed state of the protein, after furin cleavage.^18^ The S protein is a homotrimer:^21^ each monomer is shown in a different colour, namely green, cyan and orange, with glycans depicted in pink. Each monomer is formed by three domains: head, stalk and cytoplasmic tail (CT).^21^ The Y674-R685 region is shown in red. In MD simulations of the glycosylated SARS-CoV-2 S protein,^18^ Y674-R685 is accessible, being only weakly shielded by the glycans (**Figure S4**) and also shows high flexibility (**Figure S5**). (**B)** The cryoEM structure of the muscle-type receptor from *Tetronarce californica* (PDB code: 6UWZ). ^19^ This receptor is a heteropentamer formed of two α (green), one β (blue), one d (yellow), and one γ (orange) subunits. Each monomer is formed by four domains:^15-17^ extracellular (ECD), transmembrane (TMD), intracellular (ICD) and C-terminal domain (CTD). The agonist binding site is located in the ECDs at the interface between two neighbouring subunits.

According to Changeux *et al*.’s ‘nicotinic hypothesis’, direct interaction between SARS-CoV-2 and nAChRs is proposed to occur via a loop in the viral S protein^11^ (**Figure S3**). Here, we investigate this potential interaction for three nAChR subtypes, using fully solvated, all-atom molecular dynamics (MD) simulations. Our modelling (using atomically detailed structures of the glycosylated S protein,^18^ and the muscle-type nAChR from *Tetronarce californica*^19^) suggests that the association between the S protein and nAChR is not possible when the two proteins are parallel to each other but can be achieved in an orthogonal arrangement (**Figure S3**). The S protein from SARS-CoV-2 is a fusion protein^20,21^ found on the surface of the virion that mediates entry into host cells. It is an extensively glycosylated homotrimer with each monomer formed by three domains (**Figure 1A**): head, stalk and cytoplasmic tail (CT).^21^ The head comprises two subunits: S1 is responsible for binding to ACE2 of the host cell,^21^ and S2 for membrane fusion.^21^ The SARS-CoV-2 S protein contains two proteolytic cleavage sites:^21^ one (‘furin cleavage’ site) at the S1/S2 boundary thought to activate the protein^22^ and a second in the S2 subunit that releases the fusion peptide.^23^ The loop suggested by Changeux *et al*. to be directly involved in the interaction with nAChRs spans from Y674 to R685 and is located in the head region of the protein, at the interface between the S1 and S2 domains, immediately preceding the S1/S2 cleavage point^21^ (**Figure 1A**). Furin cleaves the peptide bond after R685, thus separating it from its neighbour S686 (e.g. before viral exit from the host cell).^22^ Cleavage activation of viral glycoproteins is known to be important for infectivity and virulence.^21,22^ Analysis of the dynamics of Y674-R685 in MD simulations of the full-length glycosylated SARS-CoV-2 S protein^18^ (see Supporting Information) reveals that the furin loop region is only weakly shielded by the glycans, and predominantly solvent exposed, especially when the S protein is in the closed state (**Figure S4**). In these simulations, Y674-R685 shows a different extent of conformational flexibility according to the state (open or closed) of the S protein, which might allude to a different binding propensity. This is most likely a structural consequence of the S protein’s receptor binding domain transitioning between the two states, which ultimately affects the packing of the three monomers. However, the peptide is found to adopt conformations potentially compatible with binding to nAChRs (**Figure S5**).

The Y674-R685 region contains the 4-residue PRRA insertion not present in other SARS-CoV-related coronaviruses,^24^ and includes a sequence motif homologous to several neurotoxins known to target nAChRs.^11^ In SARS-CoV-2, abrogation of the PRRA motif moderately affects virus entry into cells.^21,22^ This motif has recently been shown experimentally to interact with neuropilin-1 receptors^25^ and has also been suggested to have an affinity for T cell receptors.^26^ The high sequence similarities between Y674-R685 region and several known nAChR antagonists (**Figure S1**) suggests that this region of the SARS-CoV-2 S protein may bind to nAChRs, and could potentially act as an antagonist thus inhibiting gating.^11^ Hence, it has been postulated that nicotine may have an effect in COVID-19 by competing and interfering with this binding. Note that very recently an alternative region (G381 to K386 in the S1 subunit) in the S protein has been hypothesized to interact with nAChRs.^27^

Here, we use molecular simulation to examine the nicotinic hypothesis proposed by Changeux *et al*.^11^ of whether the SARS-CoV-2 S protein can bind stably to nAChRs via the Y674-R685 region. To test this, we built structural models for the complexes formed by the 12-residue region from the S protein (S-peptide) and the ECDs of three different nAChRs, namely the human α4β2, human α7 and muscle-like αβγδ receptor from *Tetronarce californica* (hereafter named αβγδ). These simulations build on our successful previous extensive simulations of nAChRs, which have e.g. identified a general mechanism for signal propagation in this receptor family.^28-30^

The α4β2 nAChR is the most prevalent heteromeric subtype in the brain: it is implicated in diverse processes such as cognition, mood and reward, and is necessary for nicotine addiction.^15-17,31^ The homomeric α7 nAChR is also abundant and widely expressed in the CNS, where it contributes to cognition, sensory processing and attention.^32^ The α7 subtype is also expressed on a variety of non-neuronal cells, such as immune cells, astrocytes, microglia, endothelial cells, where it contributes to anti-inflammatory pathways.^33-35^ Due to its role in the downregulation of the production of pro-inflammatory cytokines,^33-35^ it has been suggested that the α7 nAChR may be involved in the hyper-inflammation response that can be caused by SARS-CoV-2.^9,36^ The muscle-type receptor derived from the electric organ of *Tetronarce californica* (formerly *Torpedo californica*) is one the most extensively studied nAChRs, and has provided significant structural insight into this receptor family. It is formed by two α and one β, δ and γ subunits and has high sequence similarity (55%-80% identity) with its human counterpart.^37^ For this reason, and because of the availability of structural information, we used it here as a proxy for the human muscle-type nAChRs. Muscle fatigue, myalgia and arthralgia are common symptoms in COVID-19 patients, but we are not aware of studies reporting the presence of SARS-CoV-2 in the skeletal muscles, joint, or bones. It is still unclear how the virus affects the musculoskeletal system.^38^

Structural models of the three SARS-CoV-2 S-peptide–nAChR complexes were built based on the cryoEM structure of the αβγδ receptor from *Tetronarce californica* with bungarotoxin.^19^ α-bungarotoxin is a neurotoxin that acts as a nAChR antagonist, directly competing with acetylcholine,^39^ and has high sequence similarity with the Y674-R685 region of the S protein of SARS-CoV-2 (**Figure S1**). Twenty models were generated for each complex, and the one with the lowest Modeller objective function^40^ (**Figures 2 and S6**) was used as the starting point for MD simulations (see the Supporting Information for more details). Three replicates, each 300 ns long, were performed for each complex to investigate the peptide-receptor conformational behaviour and possible induced-fit effects.

**Figure 2.**
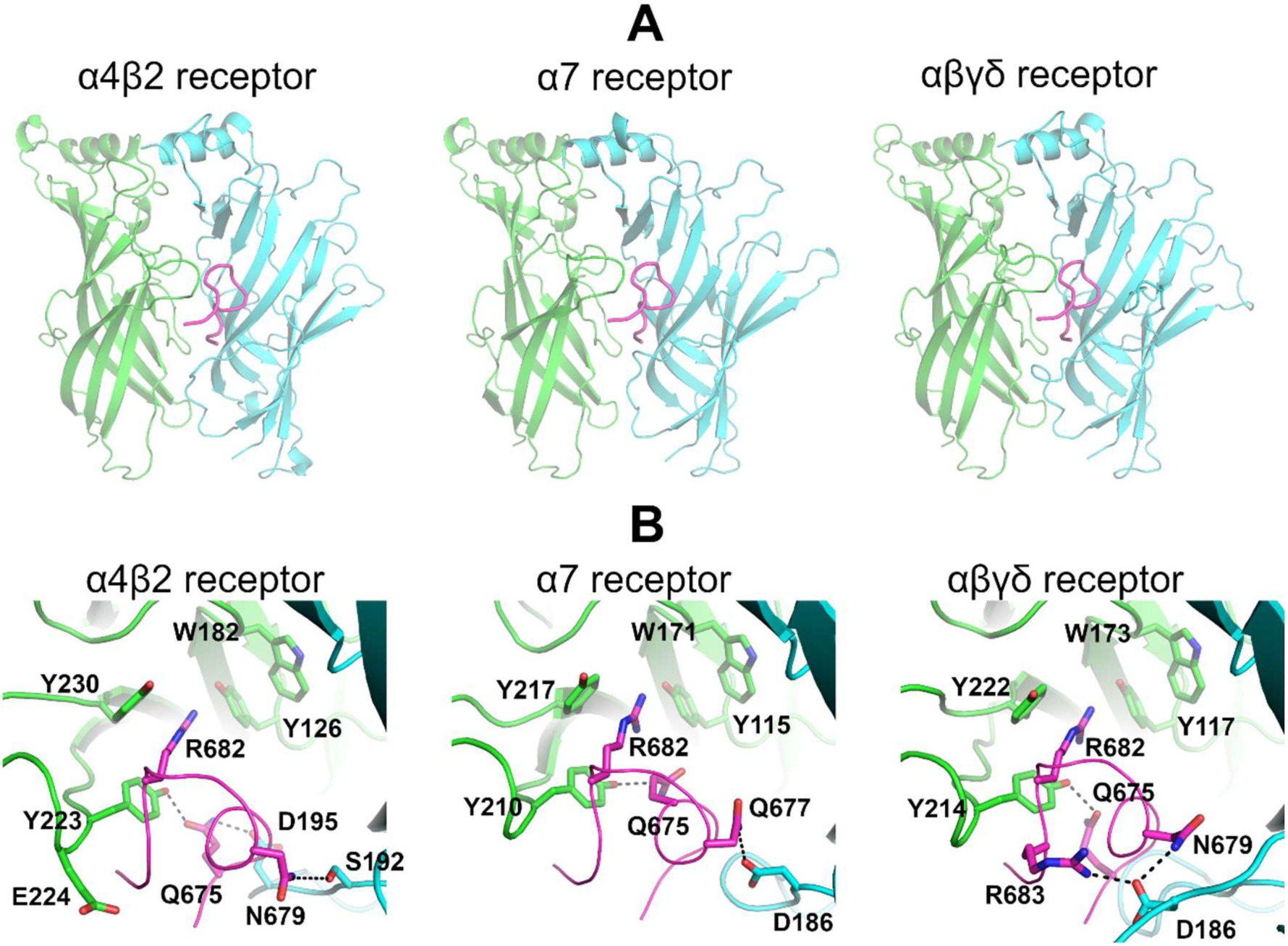
Predicted binding modes of the SARS-CoV-2 S-peptide to different nAChRs. **(A)** Complexes formed by the S-peptide and three different nAChRs, namely the human α4β2, human α7 and the muscle-like αβγδ receptor from *Tetronarce californica*. The S-peptide (region Y674-R685) is highlighted in magenta, and the principal and complementary subunits of the receptors are coloured in green and cyan, respectively. These models show the conformation of the S-peptide bound to the first pocket. In the human α4β2 receptor, the binding pocket is formed by one α4 and one β2 subunit, whereas in the human α7 nAChR, the pocket is formed by two α7 subunits. In the αβγδ receptor, the two binding pockets are non-equivalent: one is formed by an α and a δ and the second by an α and a γ subunits. (**B)** Closeup view of the peptide-receptor interaction region. Residues involved in binding of the S-peptide are shown with sticks. Note that the sidechain of R682 in the S-peptide is located inside the aromatic box establishing cation-π interactions with some of the highly conserved aromatic residues lining the pocket. Note also that all residue numbers used in this work, unless stated otherwise, refer to the human α7 (UniProt code P36544), human α4 (UniProt code P43681), human β2 (UniProt code P17787), *Tetronarce californica* α (UniProt code P02710), *Tetronarce californica* δ (UniProt code P02718), *Tetronarce californica* γ (UniProt code P02714) and SARS-CoV-2 S protein (Uniprot code P0DTC2) sequences.

At the beginning of the simulations, the S-peptide was located in the binding pocket, bound by interactions with both the principal and complementary subunits (**Figures 2 and S6**). A closeup view of the peptide-receptor interface reveals extensive contacts (**Figures 2B and S6B**), mainly with the principal subunit. In all three complexes, the sidechain of R682 of the S-peptide binds as the recognised positively charged group, a strictly conserved pharmacophore of all nAChR ligands.^41,42^ As can be seen in **Figure 2B**, the guanidinium group of R682 is well-positioned inside the aromatic box, forming several cation-π interactions with TyrC1 (α4Y223, α7Y210, αY214 in the human α4β2, human α7 and muscle-like αβγδ receptor from *Tetronarce californica*, respectively), TyrC2 (α4Y230, α7Y217, αY222) and TyrA (α4Y126, α7Y115, αY117). Note that these cation-π interactions do not entirely mimic the binding of nicotine as no interactions with TrpB are present.^43^ R682 is part of the four-residue PRRA insertion not found in other SARS-like coronaviruses,^24^ and it forms part of the furin cleavage site located the boundary between the S1 and S2 subunits.^21^ Additional binding interactions with the peptide are also observed with different residues depending on the receptor subtype: in the α4β2 nAChR, hydrogen bonds involving the sidechains of α4Y223, α4E224, β2S192, β2D195 in the receptor and Q675, N679 and the main-chain nitrogen of A684 of the S-peptide are observed; in the α7 nAChR, two hydrogen bonds between α7D186 and α7Y210 in the receptor and S-peptide Q675 and Q677 are seen; in the αβγδ receptor from *Tetronarce californica*, hydrogen bonds involving αY214 and δD186 from the receptor and Q675, N679, R682 and R683 of the peptide are observed.

During the simulations, distinct patterns of dynamical behaviour were identified for the S-peptide in the different receptor subtypes. In the α4β2 and α7 complexes, the peptide showed high positional and conformational variability, while in the αβγδ complex, it generally remained in the same pose throughout the simulation (**Figures S7 and S9**). Similar behaviour is observed for the peptides in the two binding pockets in each complex. When bound to the α4β2 and α7 nAChR, the peptide adopted many different binding modes inside the pocket, ranging from highly compact to fully extended conformations (**Figure S9**). In contrast, in the αβγδ receptor, the peptide was more compact (**Figure S9**). The range of the radius of gyration values for the S-peptide in all three complexes is similar to that observed in the simulations of the full-length glycosylated SARS-CoV-2 S protein embedded in a viral membrane^18^ (**Figures S5**). Principal component analysis (PCA) of the dynamics of the peptide revealed different conformational behaviour of the peptide in the three complexes. When bound to the muscle-like receptor, the peptide shows limited dynamical freedom: it explores a restricted conformational space spanned by the first two principal components (**Figure S10**).

The number of hydrogen bonds between the peptide and the receptors was determined over the simulations (**Figure S11**). Two more H-bonds are observed in the αβγδ complex than in the α4β2 and α7 receptors (**Figure S11**). These additional interactions with the complementary subunit (**Figure S11**) probably contribute to the increased stability of this complex and the more compact conformation of the peptide in the αβγδ receptor.

Analysis of the distribution of the distance between the R682 of the peptide and the conserved aromatic residues forming the aromatic box shows the distinctive behaviour of the peptide bound to different receptors (**Figure S12**). Interactions with R682, TyrC1 and TyrC2 are quite frequent in all three complexes, being present more than 60% of the time. To examine how deeply in the binding pocket the peptide inserts, we monitored the interactions of R682 with TrpB, a residue lining the back wall of the nAChR aromatic box. TrpB (α4W182, α7W171 and αW173) is highly conserved across the nAChR family, and it makes cation-π and H-bond interactions with the positively charged group on the ligands.^41,42^ In the α4β2 and αβγδ complexes, the S-peptide does not extend far into the pocket and interactions between R682 and TrpB are mostly absent (**Figure S12**). In contrast, in the α7 complex, the peptide binds more deeply into the hydrophobic cavity, adopting conformations that allow not only for the direct contact between R682 and TrpB (**Figures S13-S14**) but also achieve optimal core-binding interactions (**Figure 3**). In such configurations, other interactions are present in addition to those with TrpB, namely cation-π interactions with TyrC1 and TyrC2 (**Figure S14**). Although no direct contact between R682 and TyrA is observed, both residues are connected through a H-bond network mediated by Q675 from the S-peptide (**Figure S15**). This is significant because interactions with TyrA, TrpB, TyrC1 and TyrC2 are known to be critical for ligand binding and to modulate gating in the α7 subtype.^44-46^

**Figure 3.**
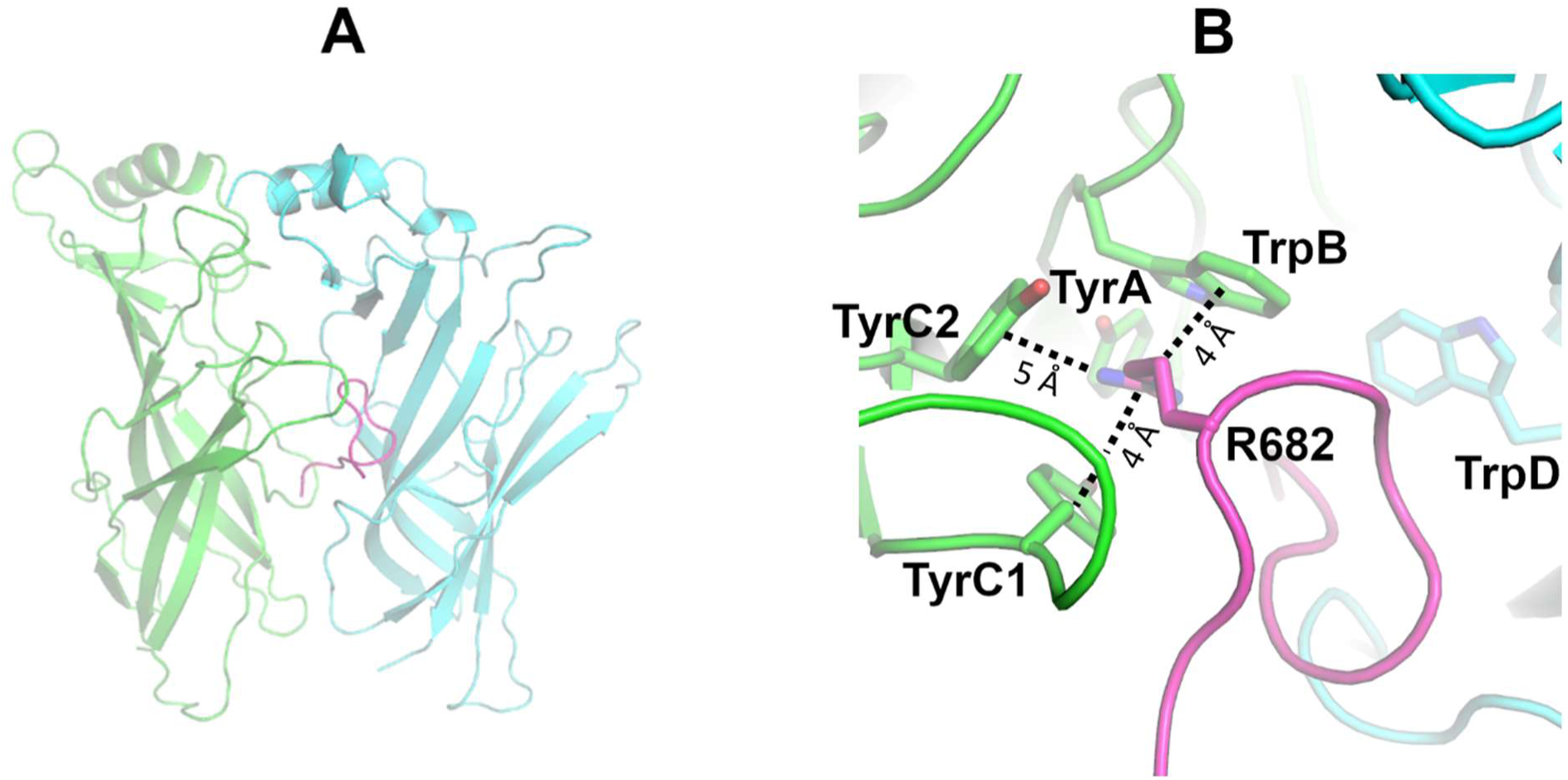
Representative conformation of the α7 complex, in which direct interaction between TrpB and R682 is observed. **(A)** Overall view of the S-peptide:α7 complex. (**B)** Closeup view of the R682 interaction region within the aromatic box. The principal and complementary subunits of the α7 receptor are coloured in green and cyan, respectively. The S-peptide is highlighted in magenta. Interactions between the guanidinium group of R682 and the aromatic rings of TrpB (α7W171), TyrC1 (α7Y210) and TyrC2 (α7Y217) are shown with dashed lines.

The binding of a ligand or a peptide can be expected to affect the conformational dynamics of the receptors. To investigate this, the Root Mean Square Fluctuations (RMSF) profiles of the C_*α*_ atoms were determined for all three receptors. Distinct dynamic behaviours are observed for the binding site regions (**Figures S16-S18**). Such differences are mostly located in loops C and F, two structural motifs important for binding and selectivity.^45,47,48^ Loop F shows decreased flexibility in the α4β2 complex, while loop C dynamics is more restricted in the muscle-like αβγδ receptor, compared to the other two subtypes.

At the beginning of the simulations, in all the three complexes, loop C adopted an open conformation due to the steric interference of the peptide. During the simulations, the αβγδ and α4β2 receptors mostly maintained this open conformation. In the α7 complex, as the peptide moves deeper into the binding pocket, loop C rotates inwards, adopting a semi-closed structure. Loop C capping is known to be important for the anchoring of the ligands into the binding pocket^45,47^ and has been suggested to be indirectly involved in gating.^29,49^ A possible relation between loop C position and ligand activation has also been proposed,^48^ with agonists stabilising more compact loop conformations and antagonists disfavouring proper loop closing. Our findings could indicate that the S-peptide may act as an antagonist in the αβγδ and α4β2 receptors whereas in the α7 receptor it is unclear whether the peptide works as an agonist or antagonist.

A molecular mechanics Poisson–Boltzmann surface area (MM-PBSA) approach^50,51^ was employed to estimate the free energy of binding of the S-peptide to the different receptors (**Table S1**). MM-PBSA calculations have emerged as an efficient and useful method to determine binding free energies,^50,51^ and are widely used to study protein-ligand interactions in medicinal chemistry,^52-54^ including in drug design for nAChRs.^55,56^ The favourable calculated binding energies suggest stable complex formation between the S-peptide and all three nAChRs (**Table S1**).

*In silico* alanine-scanning mutagenesis was performed to identify important residues (referred to as ‘hot-spots’) driving peptide–receptor association (**Figures S19-S21**). Hot-spots are residues with high energetic contributions to the thermodynamic stability of a given complex.^57^ Alanine-scanning methods provide a detailed energy map of a protein-binding interface.^57^ Here, we used the fast *in silico* method, BudeAlaScan,^57^ in which every residue, for both receptor and peptide, is mutated to alanine, and hot-spots are determined by the difference between the binding free energies of the mutant and wild-type complexes (ΔΔ*G*_bind_).^57^ Hot-spots were identified at the interface of the receptor, some of them common to all three subtypes (**Figure S22** and **Tables S2-S3**). In particular, TyrC1 (α4Y223, α7Y210, αY214) and the negatively charged residues in the upper part of loop F (β2D195, α7D186, dD201, dE203) strongly stabilise the complex. In the human α7 nAChR, the substitution of several key agonist-binding residues within the aromatic box, namely TyrA (α7Y115), TyrC1 (α7Y210), TrpB (α7W171) and TrpD (α7W77), by alanine is also predicted to destabilise the interface between the peptide and the receptor. Concerning residues in the peptide, Y674, R682 and R685 are the major contributors to stabilizing the interface (**Figure S23**). Overall, this analysis reinforces the critical role of R682 in binding to nAChRs.

In summary, the findings reported here support the hypothesis that the SARS-CoV-2 S protein can interact with nAChRs. Our calculations indicate stable binding of the S protein to these receptors through a region adjacent to the furin cleavage site and corresponding to the Y674-R685 loop. They also show apparent subtype-specific interactions and dynamics for the Y674-R685 region. COVID-19 is known to cause a range of neurological,^58,59^ muscular,^38^ and respiratory^60^ symptoms, and these predicted interactions may be relevant to understand the pathophysiology associated with this disease.

Our results indicate that the Y674-R685 region from the S protein has affinity for nAChRs in general. The region in the S protein responsible for the binding to nAChRs harbours the PRRA motif and shares high sequence similarity with neurotoxins known to be nAChRs antagonists. In particular, the guanidinium group of R682 is the key anchoring point to the binding pocket, where it forms several interactions with the residues that form the aromatic box. Analysis of the structure and dynamics of the full-length glycosylated S protein shows that the Y674-R685 region protrudes outside the glycan shield, and is flexible, showing that it is accessible to bind to nAChRs (and to other receptors such as neuropilins^25^). Modelling the interaction between the full-length S protein and nAChRs indicates that association is possible with the proteins in a non-parallel orientation to one another. It has been shown experimentally that not all S proteins protrude straight from the viral surface and a tilt angle up to 60° relative to the normal axis of the membrane is observed.^61,62^ This apparent flexibility of the S-protein would facilitate binding to host nAChRs.

In the α4β2 and αβγδ complexes, the conformational dynamics of the bound Y674-R685 peptide are compatible with the hypothesis of it acting as an antagonist: it forces the loop C to adopt an open conformation and prevents the formation of key interactions within the binding pocket. Intriguingly, in the α7 complexes, the peptide adopts binding modes that allow for the establishment of strong interactions within the aromatic box, thus raising the question of whether, in this subtype, it promotes gating. This is important because activation of α7 nAChR triggers anti-inflammatory signalling mechanisms in inflammatory cells, leading to a decrease in cytokine production, which may have relevance in understanding COVID-19 pathology. If nicotine does indeed prove to have any clinical value, it is likely that it would be due to interfering with the association with nAChRs. If so, nicotine analogues (e.g. smoking cessation agents) such as varenicline,^63^ cytisine^64^ and cytisine derivatives^30^ could also find useful application for COVID-19.

Given the promising results presented here, structural, mutational and single-channel studies will be of interest to test the importance of the interactions of the S protein from SARS-CoV-2 with nAChRs, and the potential relevance of these interactions to pathology and infectivity in COVID-19.

## Acknowledgements

AJM and ASFO thank EPSRC (grant number EP/M022609/1) and Elizabeth Blackwell Institute for Health Research, University of Bristol, for financial support (Elizabeth Blackwell Institute Rapid Response Funding Call (COVID-19). MD simulations were carried out using the computational facilities of the Advanced Computing Research Centre, University of Bristol (http://www.bris.ac.uk/acrc) and Oracle Public Cloud Infrastructure (https://cloud.oracle.com/en_US/iaas) under an award for COVID-19 research. We thank Dr Simon Bennie and Dr Jonathan Barnoud for help with the Cluster-in-the-cloud tool and the creation of a fully elastic and scalable cluster on the Oracle Cloud. AJM, ASFO, RBS and DKS also thank EPSRC for provision of ARCHER HPC time through HECBioSim (HECBioSim.ac.uk) under a COVID-19 award. RA acknowledges support from NIH (GM132826), NSF RAPID (MCB-2032054), an award from the RCSA Research Corp. and a UC San Diego Moore’s Cancer Center 2020 SARS-COV-2 seed grant. LC is funded by a Visible Molecular Cell Consortium fellowship. RA, LC and ZG thank the Texas Advanced Computing Center (TACC) Frontera team and acknowledge computer time made available through a Director’s Discretionary Allocation (made possible by the NSF award OAC-1818253).

