## Supplementary Information for "Simulations support the interaction of the SARS-CoV-2 spike protein with nicotinic acetylcholine receptors"

### Materials and methods

#### Modelling the interaction between nAChR and Y674-R685 from S protein

The structural models for the complexes formed by the extracellular domains of the human α7, human α4β2 and muscle-like αβγδ nAChR from *Tetronarce californica* and the Y674-R685 region from SARS-CoV-2 (hereafter named as S-peptide) were constructed using the cryoEM structure of the αβγδ receptor from *Tetronarce californica* (formerly *Torpedo californica*) with α-bungarotoxin[1] (PDB code: 6UWZ) as a template. Note that the homology model for the ECDs of human α7 nAChR was constructed because there is no Xray or cryoEM structure for this nAChR sub-type in the Protein Data Bank. Note also that the experimentally-determined structures available for the human α4β2 nAChR[2,3] have nicotine bound in the binding pockets and as such show loop C in the  a closed ‘‘capped’’ conformation. Loop C acts as a binding-pocket lid and it adapts its shape to the size of the ligands.[4,5] Given that there is no experimental structure for this sub-type with loop C in the open “uncapped” conformation, a homology model was built for the ECDs of human α4β2 nAChR based in the cryoEM structure of the αβγδ receptor with an antagonist (α-bungarotoxin) bound.[1]

The structure used here as template reflects the closed state of the muscle-type αβγδ nAChR stabilized by the binding of the two α-bungarotoxin molecules at the α-γ and α-δ interfaces.[1] α-bungarotoxin is a 74-residue neurotoxin that binds to the muscle receptors in an (almost) irreversibly way[6], and it acts as nAChRs antagonist directly competing with acetylcholine.[4,5] The binding of α-bungarotoxin to the neuromuscular junction receptors induces paralysis, respiratory failure, and eventually death.[7] The sequence alignment between the S-peptide and α-bungarotoxin was taken from the work of Changeux *et al*.[8] The sequences for the different nAChRs subunits were obtained from the UniProt database:[9] human α7 (UniProt code P36544), human α4 (UniProt code P43681), human β2 (UniProt code P17787) and aligned with the template using Clustal Omega.[10,11] Twenty models were generated for each complex using Modeller 9v20.[12,13] The best model for each complex (the one with the lowest value for Modeller’s objective function[13]) was further analyzed using Procheck.[14] Overall, the α4β2 and α7 models are similar to the structures used by us in previous work,[15,16] with the exception of the loop C region that shows an open “uncapped” conformation and loop F that was slightly displaced to accommodate the S-peptide.

#### MD simulations

The best model for each complex was used as the starting point for molecular dynamics (MD) simulations. Three systems were prepared, the human α7, human α4β2 and αβγδ nAChR from *Tetronarce californica,* each with two SARS-CoV-2 S-peptides bound, one in each nonconsecutive binding pocket. The protonation state of each titrable residue in the receptor and peptides at pH 7.0 was determined using PROPKA.[17,18] All systems were solvated using TIP3P water molecules,[19] and an ionic concentration of 0.1 M sodium chloride was used. The Amber ff99SB-ILDN[20] force-field was used to describe the receptors and the peptides. All simulations were carried out in the isothermal–isobaric (NPT) ensemble at 310 K and 1 atm. The velocity-rescaling thermostat[21] and the Parrinello-Rahman barostat[22,23] were applied to keep the temperature and pressure constant. A time step of 2 fs was used for integrating the equations of motion. Non-bonded long-range electrostatic interactions were calculated using PME.[24] A 12 Å cut-off was used for the van der Waals interactions with long-range dispersion corrections for the energy and pressure.[25] The neighbour list was updated every 20 steps. The solvated complexes systems were energy minimised, equilibrated (for 1.5 ns) and simulated using the protocol described in our previous work.[16] Three unrestrained MD simulations, each 300ns long, were performed for each complex. All equilibrium MD simulations were performed using Gromacs 2019[26] on the University of Bristol's High-Performance Clusters (BlueCrystal4 and BluePebble) and the Oracle Cloud Infrastructure (https://cloud.oracle.com/en_US/iaas). Accompanying preliminary simulations of the S protein were run on ARCHER using time provided by EPSRC through HECBioSim under a COVID-19 call. These were based directly on the previous work of Casalino et al.[27]

#### Analysis

Analyses were performed using Gromacs[26] and in-house tools. Images were produced with PyMOL.[28,29] Principal component analysis (PCA) was used to examine the sampling of the peptide and to identify its relevant motions. All replicates for the three complexes were combined before the analysis so that all share the same subspace, and their motions could be directly compared. 5400 frames (corresponding to one conformation per nanosecond per replicate per peptide) were used for the PCA. The two principal components included ~53% of the peptide dynamics and, hence, we restricted our analysis to PC1 and PC2 only.

PCA was also used to assess the sampling and equilibration/relaxation of the receptors similarly to e.g.[30,31] For this, all replicates for each complex were combined, and two conformations per nanosecond per replicate (totalling 1801 frames) were used for this analysis. PCA, together with the RMSD of the receptors over time, suggests that all systems were equilibrated after 50 ns (**Figure S8**). The RMSD values for the ECDs of the human α4β2 and α7 nAChR are consistent with our previously published simulations.[15]

#### MM-PBSA calculations

A Molecular Mechanics Poisson–Boltzmann Surface Area (MM-PBSA) approach was used to calculate the binding free energy ($\Delta G_{\mathrm{bind}}$) for each complex. In this approach, the contribution of nonpolar, polar and entropic terms to the overall free energy of binding is estimated from a MD simulation of the solvated complex.[32,33] Snapshots were taken every two nanoseconds per replicate per complex (in a total in a total of 453 frames per complex). Binding free energies were computed using g_mmpbsa.[34] This tool uses Gromacs[26] and APBS[35] to determine the binding energy and energetic contribution of each residue. The solvent-accessible surface area (SASA) model was used to calculate the non-electrostatic contribution to the solvation free energy, whereas the electrostatic contribution was estimated by solving the Poisson−Boltzmann (PB) equation. For the APBS calculations, a grid spacing of 0.5 Å was used with a twofold expansion in each dimension. An ionic strength of 0.10 M was used with radii of 0.95 and 1.81 Å for sodium and chloride ions. The entropy change on binding is particularly challenging to compute for the binding of a long, flexible peptide and shows high standard error compared to the other energetic terms. The entropic contribution will disfavour binding in all three receptors and is likely to be similar in all three. The calculated values are therefore most usefully analysed in terms of relative binding affinity to the three receptors.

##### *In silico* alanine-scanning mutagenesis

*In silico* alanine-scanning mutagenesis involves the sequential mutation of the residues in the proteins to alanine to identify the key determinants for the thermodynamic stability of a given complex. In this approach, the binding free energies for the mutant and wild-type complexes are calculated, and the difference between the two values ($\Delta\Delta G_{\text{bind}}$) is a way to evaluate the contribution of each reside for the interface. In this work, the $\Delta\Delta G_{\text{bind}}$ was computed using the command-line Python application BudeAlaScan.[36] This application uses ISAMBARD[37] for structure manipulation and a customized version of the Bristol University Docking Engine (BUDE)[38] for energy calculations. Snapshots were taken every three nanoseconds in a total of 303 frames per complex.

##### Molecular characterization of the S-peptide in MD simulations of the full-length model of the glycosylated S protein from SARS-CoV-2

To examine the conformational dynamics and the accessibility of S-peptide (Y674-R685 region) in the glycosylated SARS-CoV-2 S protein, we used the extensive all-atom MD simulations conducted previously by some of us (Casalino *et al*).[27] In that work, two sets of simulations were performed on two full-length models of the glycosylated S protein accounting for ~4.2 μs in the open (1 RBD 'up', 2 'down') and ~1.7 μs in the closed (3 RBDs 'down') states, which were based on 6VSB[39] and 6VXX[40] cryoEM structures, respectively. In these simulations, the models were cleaved at the S1/S2 site (i.e., between R685 and S686) to model the physiological state of S-peptide. Considering that the S protein is a homotrimer, these simulations have accumulated a comprehensive total of ~12.6 μs and ~5.1 μs of sampling for S-peptide in the open and closed systems, respectively. With the aim of elucidating the availability of S-peptide in the glycosylated full-length model of the S protein for binding to nAChRs, we have investigated the conformational behaviour of the S-peptide, and characterized its accessibility in the presence of the glycan shield.

**Accessible Surface Area**

During the simulations of the glycosylated S ptotein, the S-peptide establishes intermittent interactions with nearby N-glycans, especially N-603, N-657, N-717, N-801 and N-1074. To characterize the extent of the glycan shield, we calculated the accessible surface area (ASA) of the S-peptide (with and without glycans) using 15 different probes increasing in radius size from 1.4 Å to 15 Å, as described in Casalino *et al*.[27] Using a continuous range of values allows us to approximate different size molecules, ranging from small molecules at 2–5 Å to larger peptide- and antibody-sized molecules at 10–15 Å.[27] The ASA of the S-peptide was calculated across the replicate simulations at 2 ns intervals (**Figure S4**). Note that the difference between the overall accessibility of the 'naked' protein (without glycans) and the glycan shielded area corresponds to the effective accessibility of the S-peptide in the presence of glycans (cyan coloured area in **Figure S4**). The peptide shows a different amount of glycan shield, with an average maximum coverage (across replicas and chains) of 47% in the open system and 30% in the closed system, at 15 Å probe radii (**Figure S4**). Although the calculated ASA and glycan shield values show a marked variability due to the high flexibility of the peptide and the glycans, this analysis reveals that the S-peptide is weakly shielded, especially when the S protein is in the closed state, and potentially available for engaging with nAChRs. Interestingly, the presence of one RBD in the "up" conformation within chain A of the open system slightly alters the packing of the three monomers with respect to the closed system. This most likely results in the observed differences of accessibility and glycan shield between the two systems and even across chains (**Figure S4**). We also note that the same region of the S protein has been experimentally shown to bind to human neuropilin receptors,[41] which is clear evidence of its accessibility and ability to bind.

**Radius of gyration**

The radius of gyration (*Rg*) of the S-peptide was calculated from MD simulations of the glycosylated full-length S protein (**Figure S5**). Interestingly, the closed system exhibits a single *Rg* population, with an average of 0.78 +/- 0.09 nm, while the open system shows a wider distribution, with an average *Rg* value of 0.75 +/- 0.13 nm. Overall, the range of *Rg* values is comparable to that observed in the peptide-only simulations (**Figure S9**) and are compatible with an extended, solvent accessible conformation of the peptide (**Figure S5B**). These results further attest the different behaviour of the S-peptide between the open and closed systems possibly resulting from a slightly altered packaging of the three monomers, as also emerged from the ASA analysis. This suggests that the closed and open states may have different binding propensities.

### Supporting figures

```
SARS-CoV-2 S       674 - YQTQTNSPRRAR - 685
α-bungarotoxin      50 - CDAFCSS-RGKV -  60
RABV G             208 - CDIFTNS-RGKR - 218
```

**Figure S1.** Sequence alignment of the Y674-R685 region in the S protein from SARS-CoV-2 and two known nAChR antagonists, namely α-bungarotoxin from *Bungarus multicinctus*[4-6] and glycoprotein (G) from *Rabies lyssavirus* (formerly *Rabies virus*).[42-44] The residue numbers refer to the following UniProt codes: P0DTC2 (S protein), P60615 (α-bungarotoxin) and P15199 (G protein).

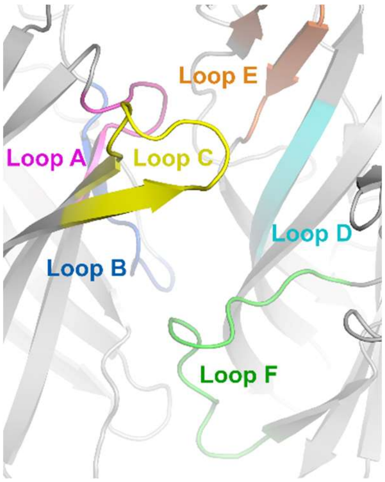

**Figure S2.** Detailed view of the nAChR binding pocket. For this image, the cryoEM structure of the muscle-type receptor from *Tetronarce californica* (PDB code: 6UWZ)[1] was used. The structural motifs lining the pocket are highlighted with the following colour scheme: loop A, magenta; loop B, blue; loop C, yellow; loop D, cyan; loop E, orange; loop E, green.

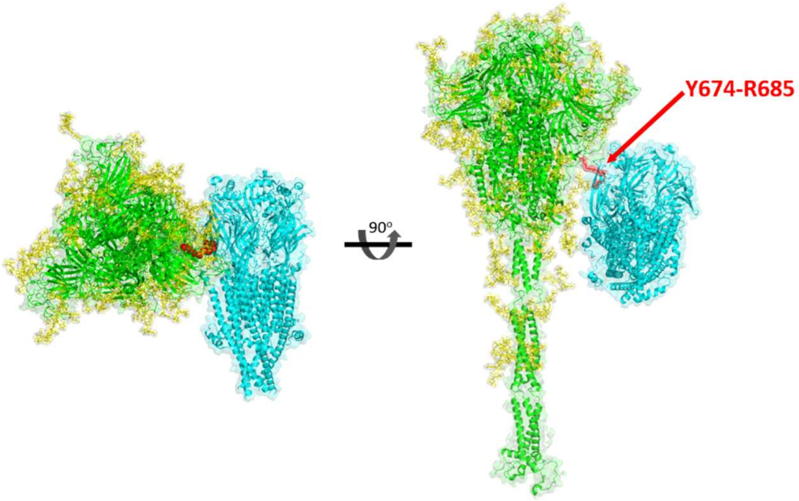

**Figure S3.** A model for the direct interaction between the SARS-CoV-2 S protein and nAChR. In this figure, the model for the full-length closed glycosylated S protein after furin cleavage was developed by Amaro and co-workers[27] whereas the for the nAChR, the cryoEM structure of the muscle-type receptor from *Tetronarce californica*[1] was used. The S protein is coloured in green (with the glycans in yellow) and the nAChR is highlighted in cyan. The Y674-R685 region in the S protein is shown with red spheres.

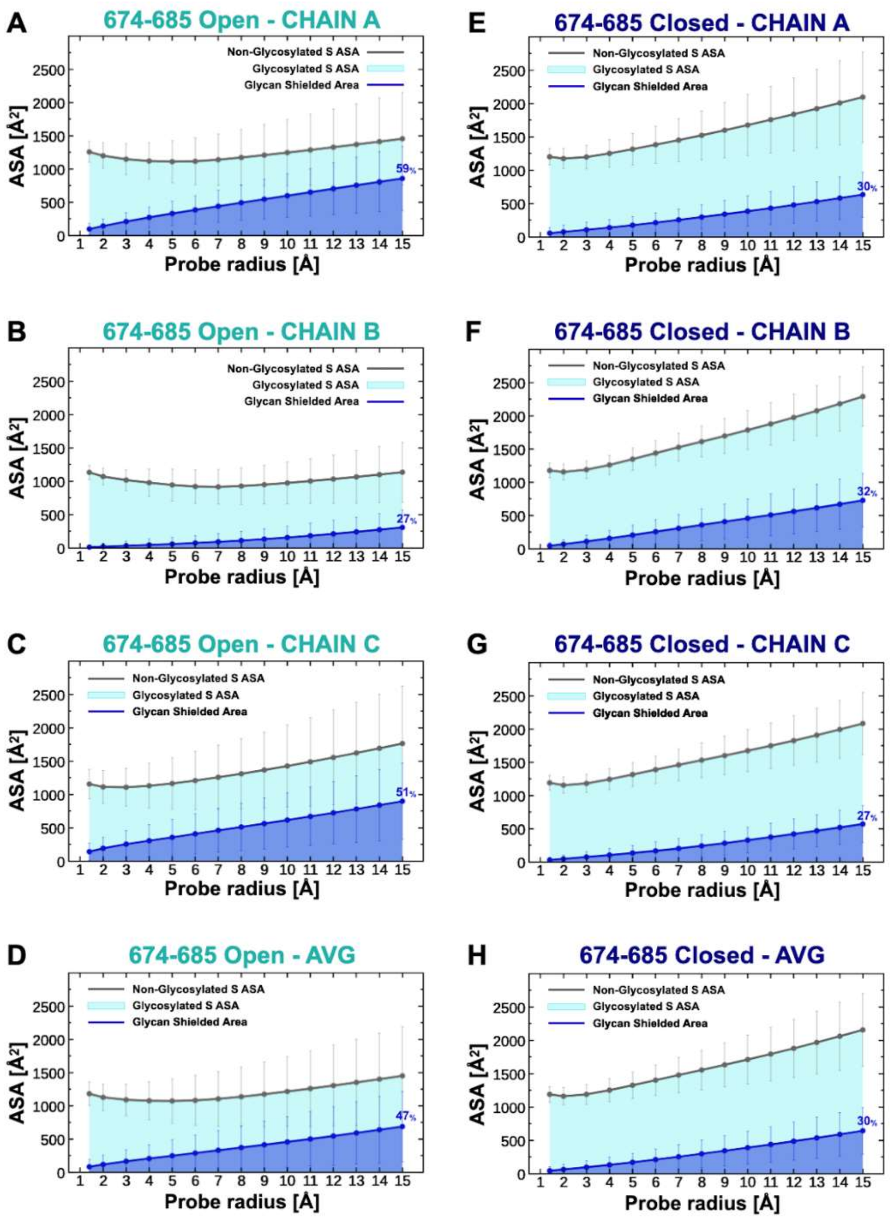

**Figure S4.** Accessible surface area (ASA) of the S-peptide in the simulations of the full-length glycosylated S protein from SARS-CoV-2.[27] The ASA of the S-peptide (residues 674-685), and the area shielded by glycans, at multiple probe radii from 1.4 (water molecule) to 15 Å are shown for the all-atom MD of the full-length models of the glycosylated SARS-CoV-2 S protein[27] in the open (**A-D**) and closed states (**E-H**). The area shielded by the glycans is presented in blue (rounded % values are reported), whereas the grey line represents the accessible area of the protein in the absence of glycans. Highlighted in cyan is the area that remains accessible in the presence of glycans. A per-chain analysis for the open state is reported in panels **A**, **B** and **C**, showing the values for the S-peptide in chain A (RBD-up), B and C, respectively. Similarly, panels **E, F** and **G** display the per-chain analysis for the closed system, where chains A, B and C are all in the 'down' conformation. The calculated values have been averaged across replicas and the error bars correspond to +/- standard deviation. Finally, in panels **D** (open) and **H** (closed), the ASA and glycans shield values were averaged also across chains.

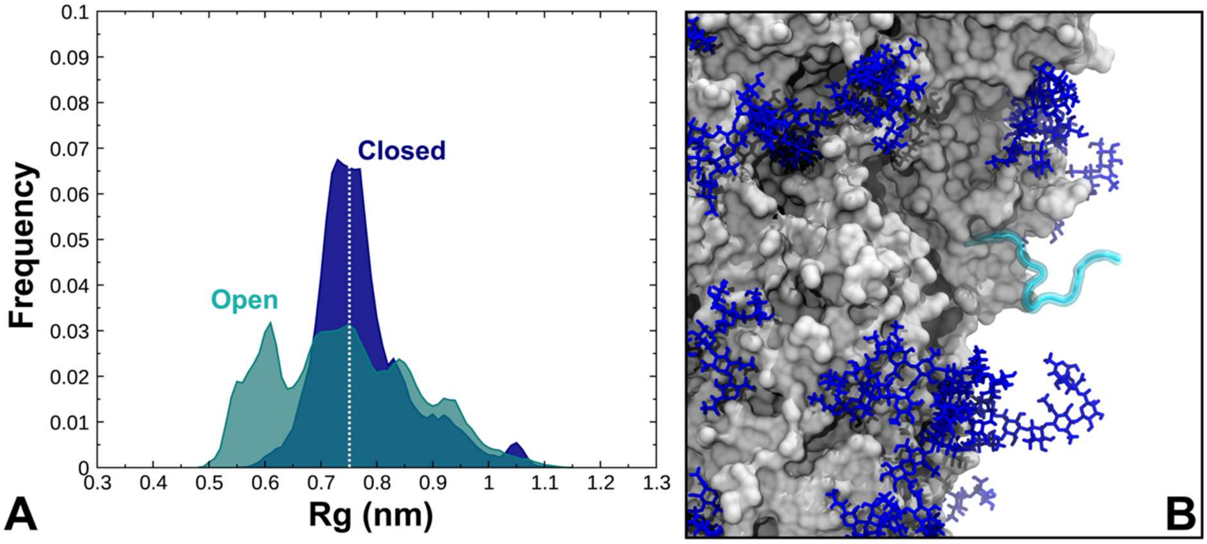

**Figure S5. (A)** Radius of gyration (*Rg*, nm) distribution for the S-peptide from simulations of the full-length model of the cleaved, glycosylated S protein in the open (teal) and closed states (blue). The two distributions are independently normalized using the respective number of data points. (**B**) A snapshot taken from the simulations of the S protein in the closed state showing one of the three S-peptides protruding into the solvent with a *Rg* of 0.75 nm. The protein is depicted with a grey surface, whereas the S-peptide is shown as a cyan ribbon. The glycans are illustrated with blue sticks.

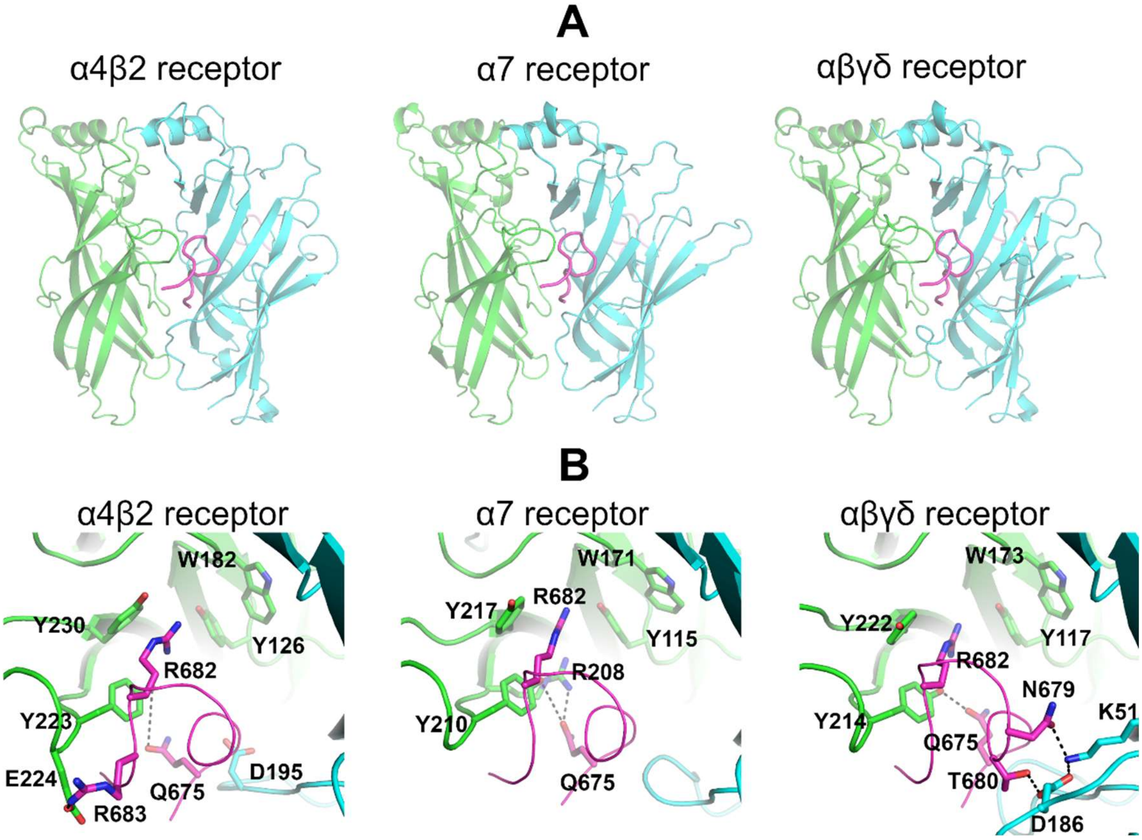

**Figure S6.** Binding of SARS-CoV-2 S-peptide to the second binding pocket in different nAChRs. **(A)** Overall view of the peptide-receptor complexes. The S-peptide (region Y674-R685) is highlighted in magenta, whereas the principal and complementary subunits are coloured in green and cyan, respectively. All three models show the peptide conformation when bound to the second pocket. (**B)** Closeup view of the peptide-receptor interaction region in each nAChR. Residues that interact directly with the peptide are shown with sticks.

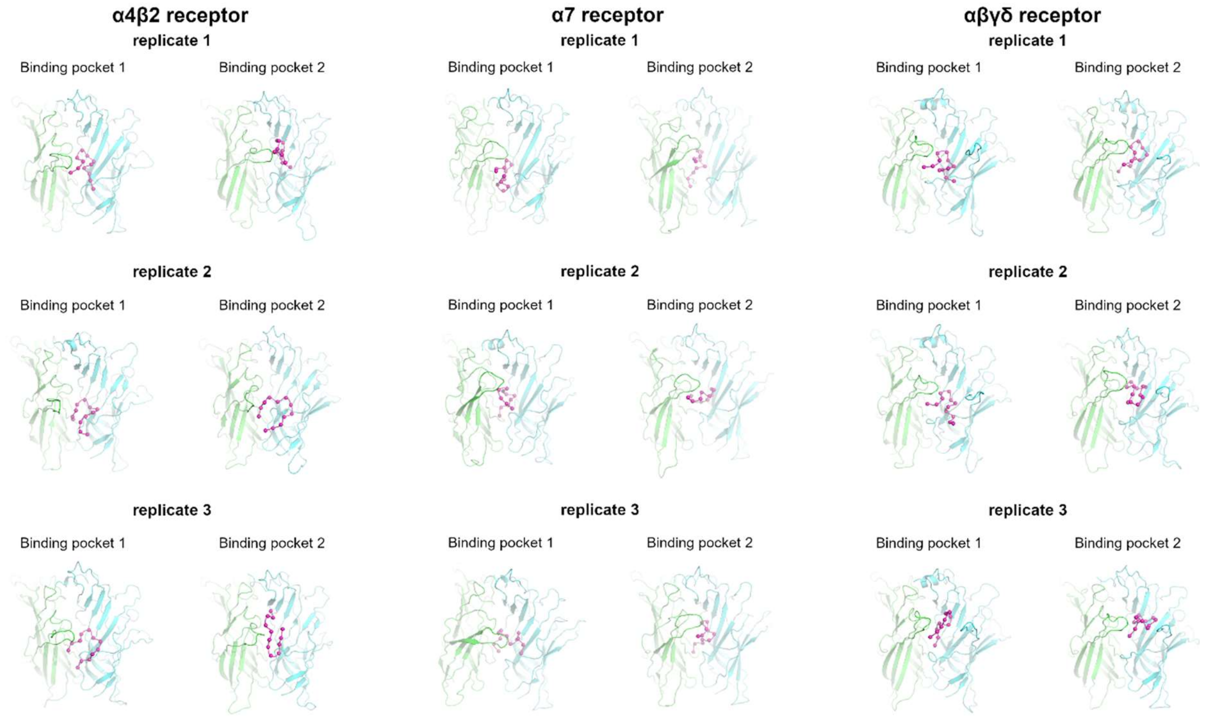

**Figure S7.** Binding mode of the S-peptide in the human α4β2, human α7 and muscle-like αβγδ receptor from *Tetronarce californica* after 300 ns of simulation, from three replicates in each case. The peptide is shown in magenta, and the principal and complementary subunits are coloured in green and cyan, respectively. Please zoom into the image for detailed visualisation.

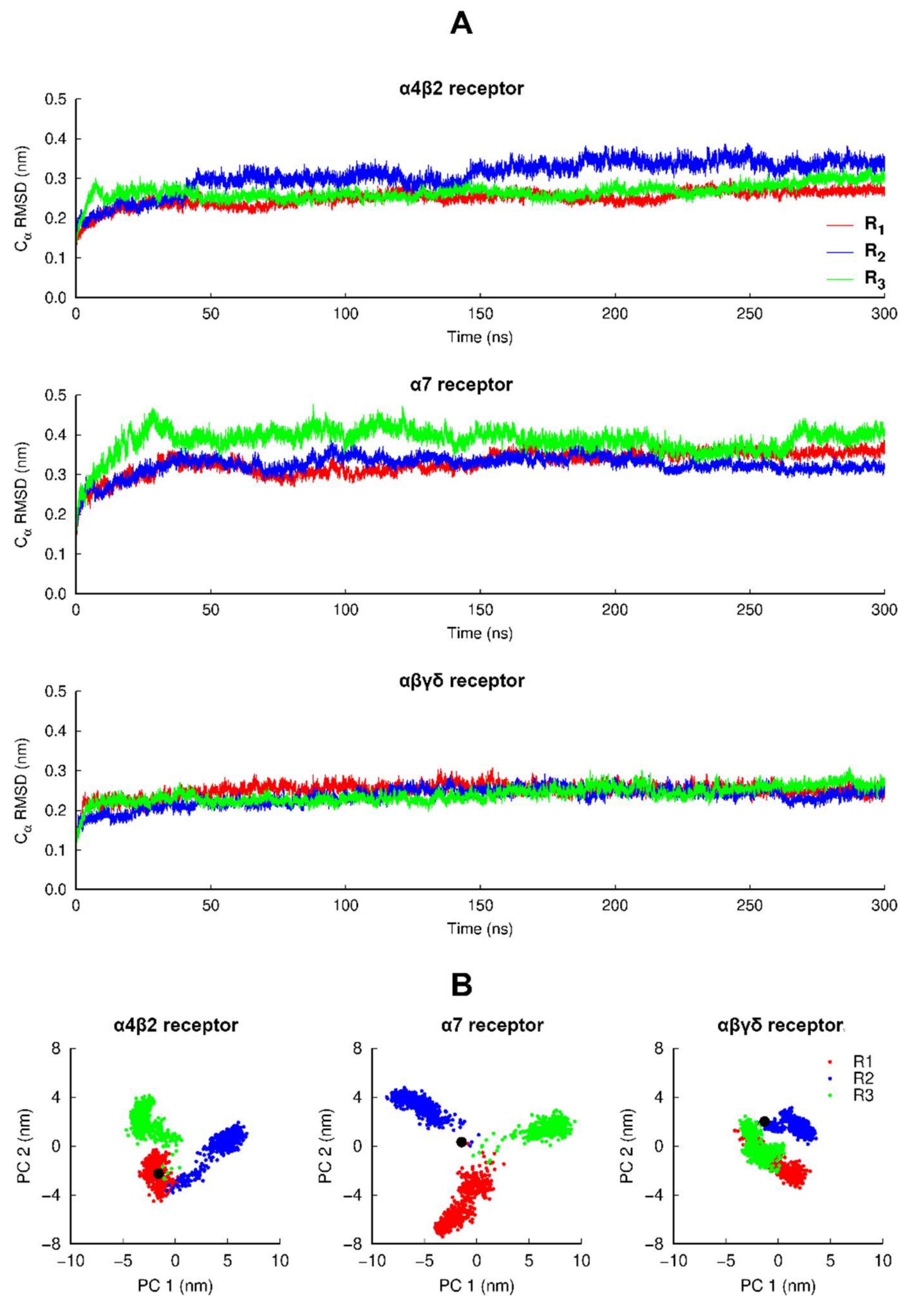

**Figure S8.** **(A)** Time evolution of the $C_\alpha$ RMSD of the individual replicates for the human α4β2, human α7 and muscle-like αβγδ receptor from *Tetronarce californica*. The $C_\alpha$ RMSD was calculated relative to the starting structures. **(B)** PCA of all replicates for the α4β2, α7 and αβγδ receptors. All three replicates for each system were combined before the analysis, and each trajectory contained two conformations per nanosecond per replicate (totalling 1801 frames) with all the $C_\alpha$ atoms of the protein. The black dot corresponds to the structure used as the starting point for the replicates. Note that the different replicates sample different regions of conformational space, thus improving the overall sampling for each system.

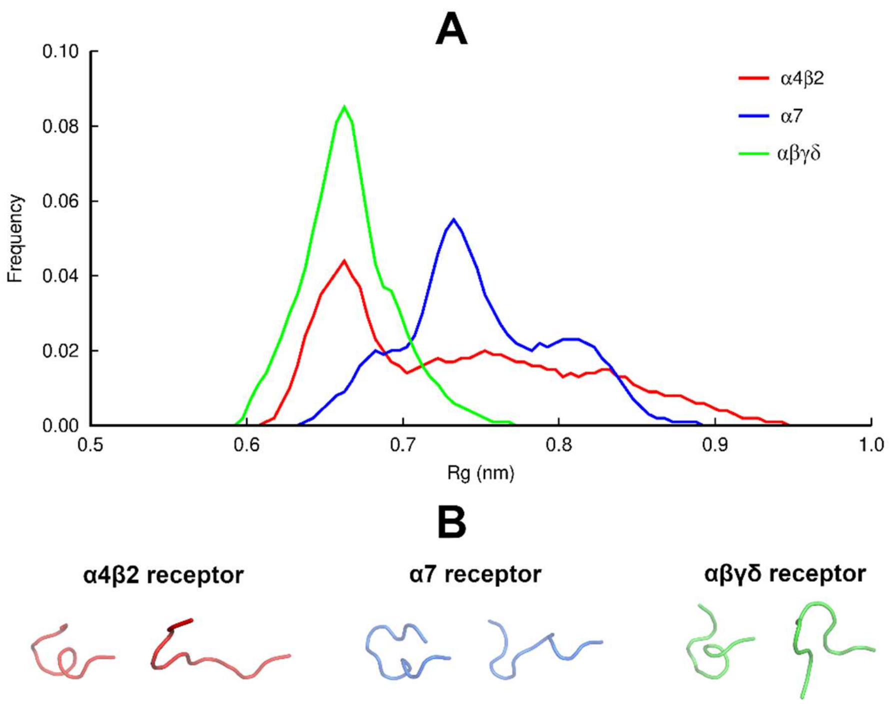

**Figure S9**. (**A)** Radius of gyration (*Rg*) distribution for the S-peptide when bound to human α4β2 (red line), human α7 (blue line) and muscle-like αβγδ receptor from *Tetronarce californica* (green line) nAChRs. The histograms reflect the *Rg* of both peptides from the 3 independent simulations performed for each complex. (**B)** The most compact and extended conformations adopted by the S-peptide when bound to the different receptors.

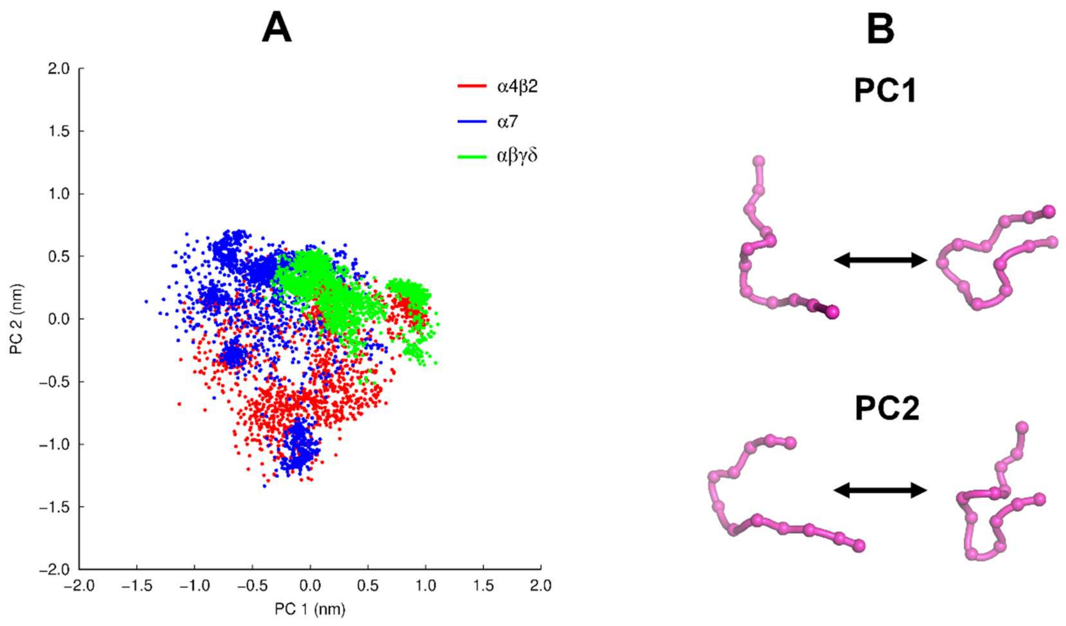

**Figure S10.** (**A)** PCA for the S-peptide (**B)** Conformations indicating motions associated with PC1 and PC2. Each trajectory contained two conformations per nanosecond per replicate per peptide (totalling 5400 frames) with all the $C_\alpha$ atoms of the peptide. PC1 and PC2 correspond to 29% and 24% of the data, respectively.

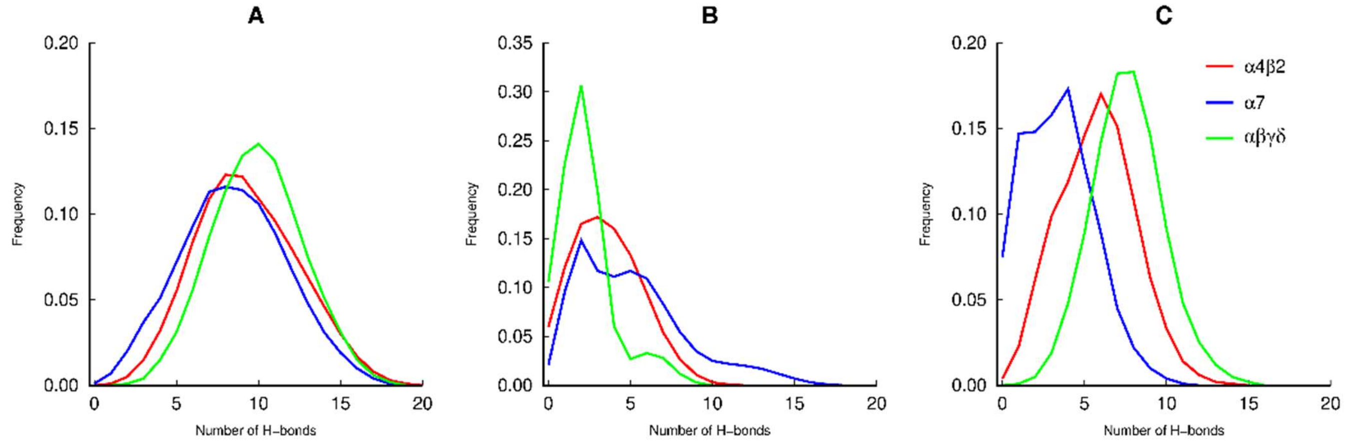

**Figure S11**. Number of hydrogen bonds between the S-peptide and the receptor. (**A)** Overall number of hydrogen bonds. (**B** and **C)** Number of hydrogen bonds with the principal (**B**) and complementary (**C**) subunits. Please zoom into the image for detailed visualisation.

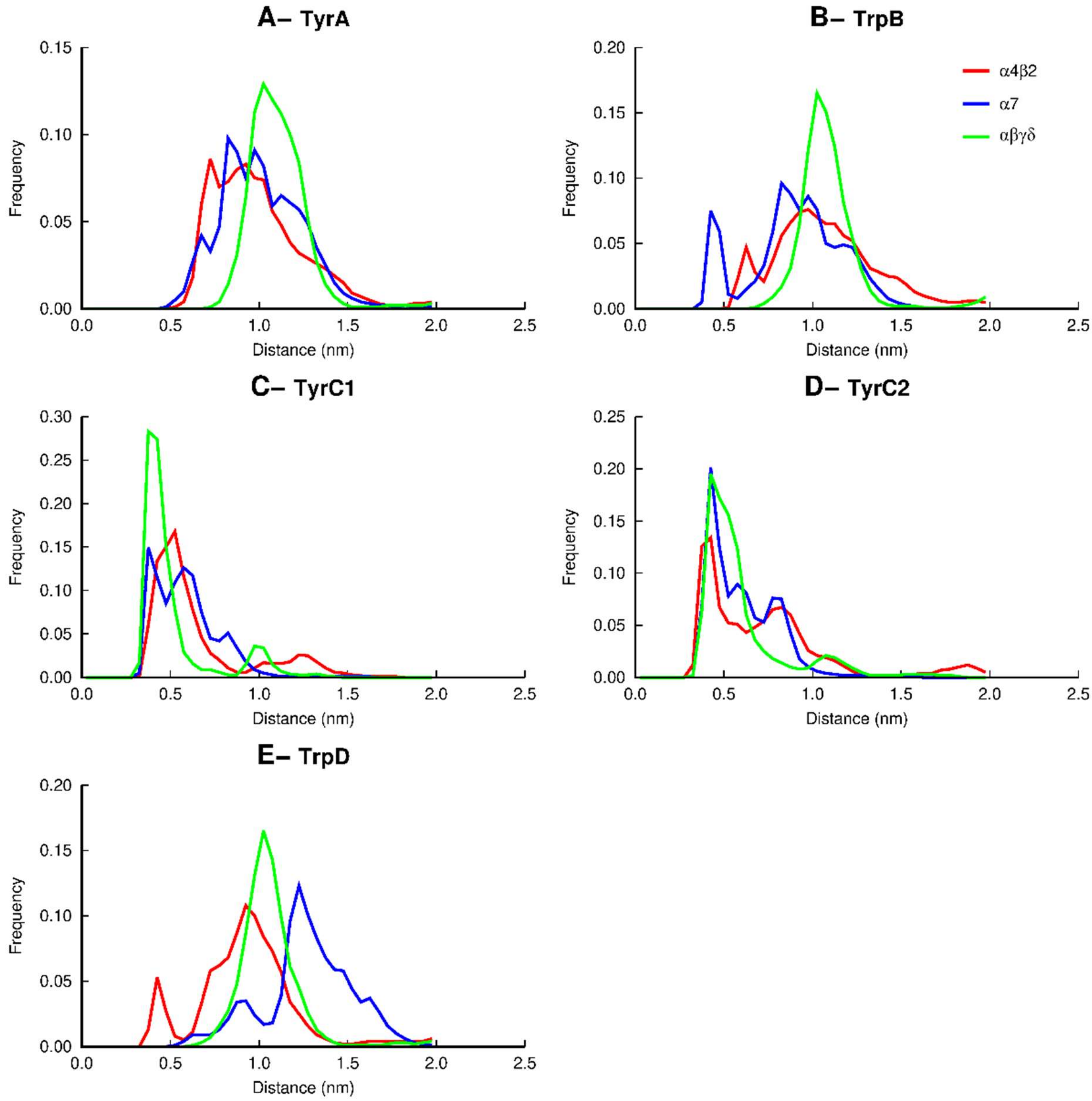

**Figure S12.** Distribution of the distance between R682 on the S-peptide and the main conserved aromatic residues lining the binding pocket. Overall distribution of the distance between the side-chains of R682 on the S-peptide and TyrA (α4Y126, α7Y115 and αY117 in the principal subunit of the human α4β2, human α7 and muscle-like receptor from *Tetronarce californica*, respectively), TrpB (α4W182, α7W171 and αW173), TyrC1 (α4Y223, α7Y210 and αY214 in the principal subunit), TyrC2 (α4Y230, α7Y217 and αY222) and TrpD (β282, α7W77, δW78 and γW72) in the complementary subunits). Note that sequence number used here refer to the following sequences: human α7 (UniProt code P36544), human α4 (UniProt code P43681), human β2 (UniProt code P17787), *Tetronarce californica* α (UniProt code P02710), *Tetronarce californica* δ (UniProt code P02718), *Tetronarce californica* γ (UniProt code P02714) and SARS-CoV-2 S protein (Uniprot code P0DTC2). The histograms reflect the distances over the two binding pockets.

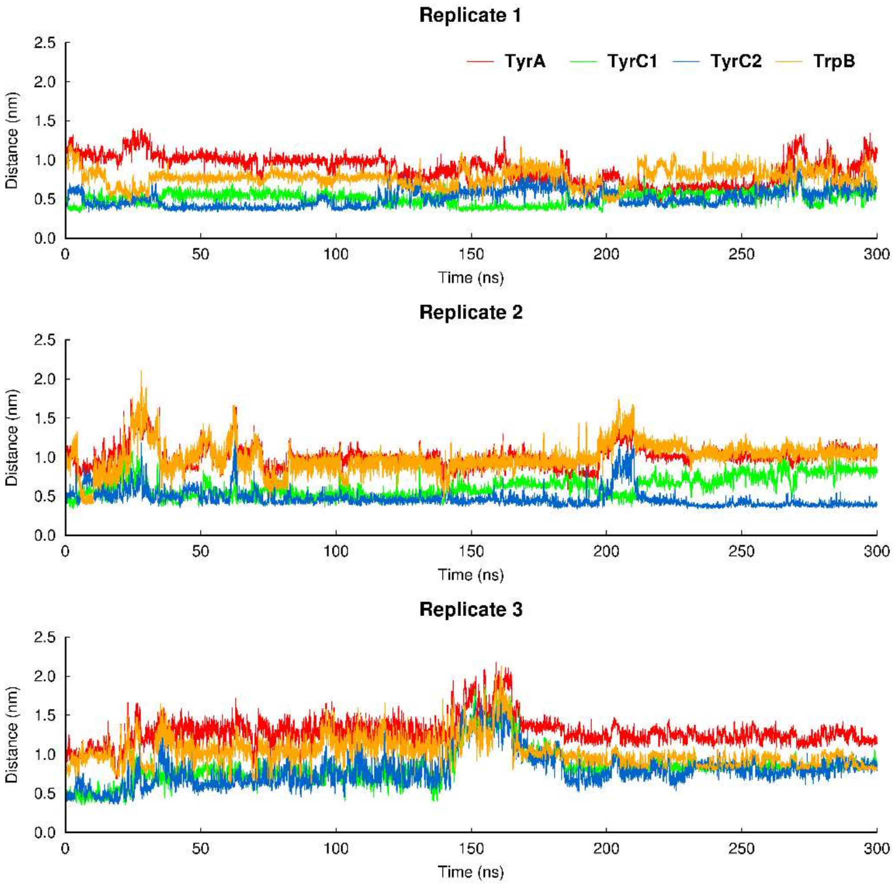

**Figure S13.** Time evolution of the distances between the sidechains of R682 on the S-peptide and TyrA, TrpB, TyrC1 and TyrC2 in the first binding pocket of the α7 nAChR.

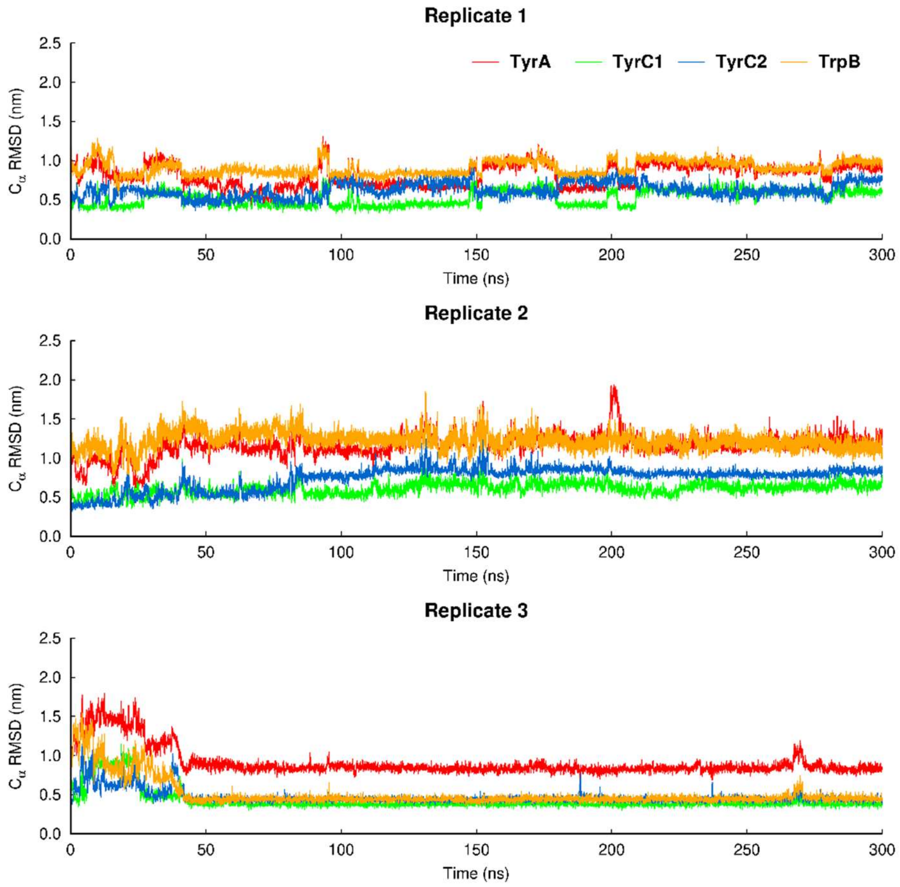

**Figure S14.** Time evolution of the distances between the sidechains of R682 on the S-peptide and TyrA, TrpB, TyrC1 and TyrC2 in the second binding pocket of the α7 nAChR.

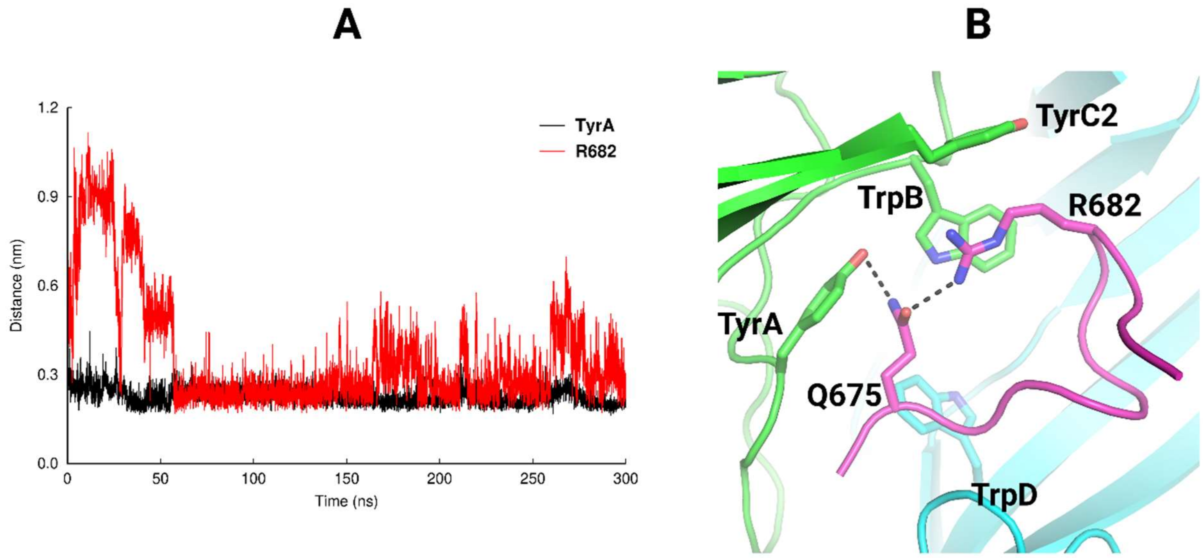

**Figure S15.** Hydrogen bond network involving TyrA in the α7 complex in which a direct interaction between TrpB and R682 is present. **(A)** Time evolution of the minimum distance between Q675 and R682 and Q675 and TyrA in replicate 3. (**B)** Closeup view of the hydrogen bond network involving TyrA, Q675 and R682 in a representative conformation of the α7 complex, in which direct interaction between TrpB and R682 is observed**.** Note that this image shows the same conformation as **Figure 3** but in a different orientation. The principal and complementary subunits of the human α7 receptor are coloured in green and cyan, respectively. The S-peptide is highlighted in magenta. Hydrogen bonds involving Q675 are highlighted with dashed lines. Part of the loop C region from the receptor was removed for clarity.

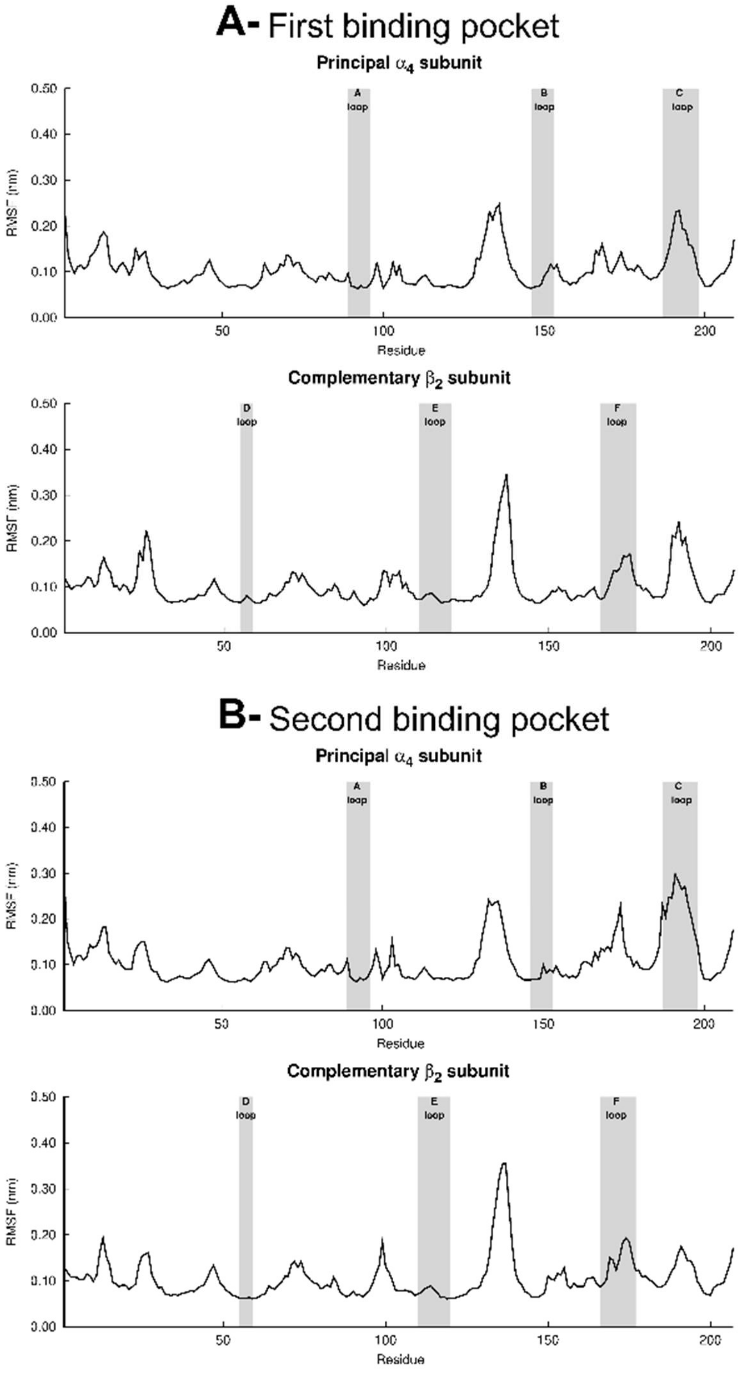

**Figure S16.** Average RMSF (averaged over the three replicates individual RMSFs) for the first (**A**) and second (**B**) binding pockets in the α4β2 complex. Please zoom into the image for detailed visualisation.

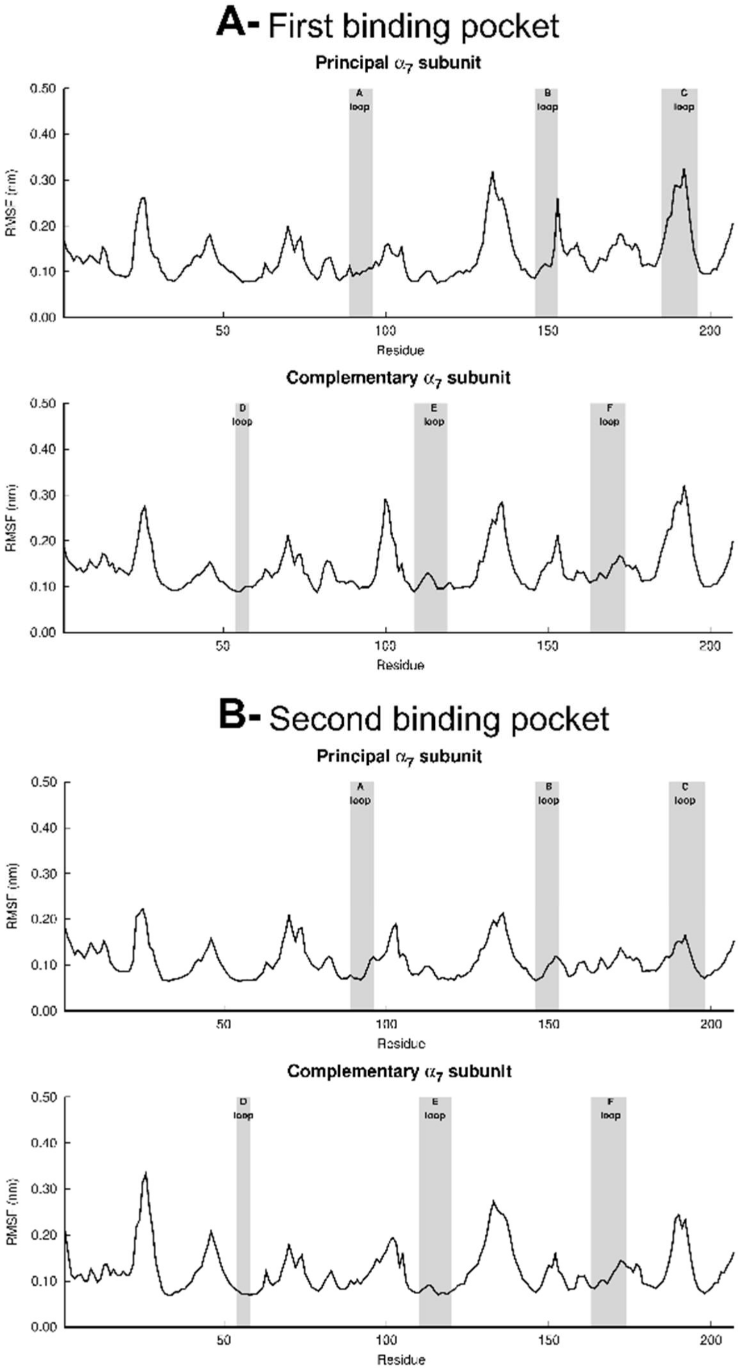

**Figure S17.** Average RMSF (averaged over the three replicates individual RMSFs) for the first (**A**) and second (**B**) binding pockets in the α7 complex. Please zoom into the image for detailed visualisation.

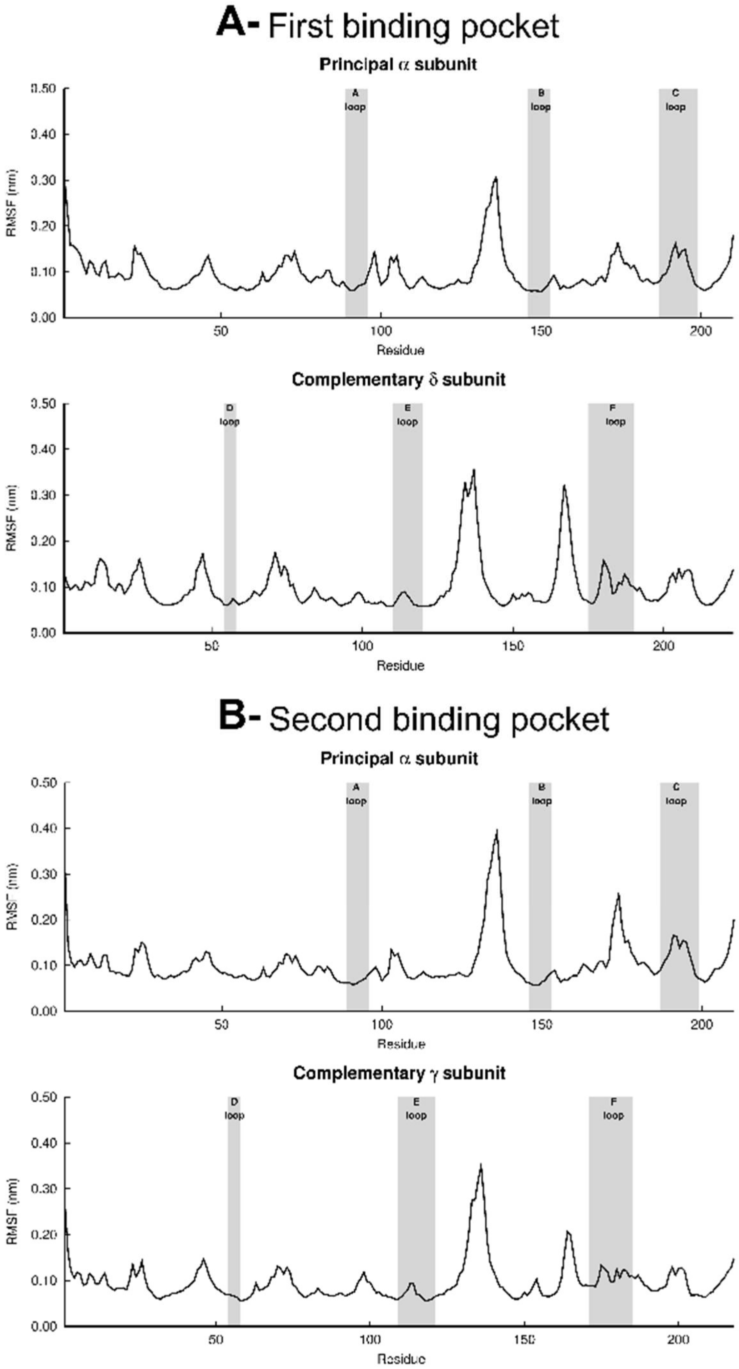

**Figure S18.** Average RMSF (averaged over the three replicates individual RMSFs) for the first (**A**) and second (**B**) binding pockets in the muscle-like αβγδ complex. Please zoom into the image for detailed visualisation.

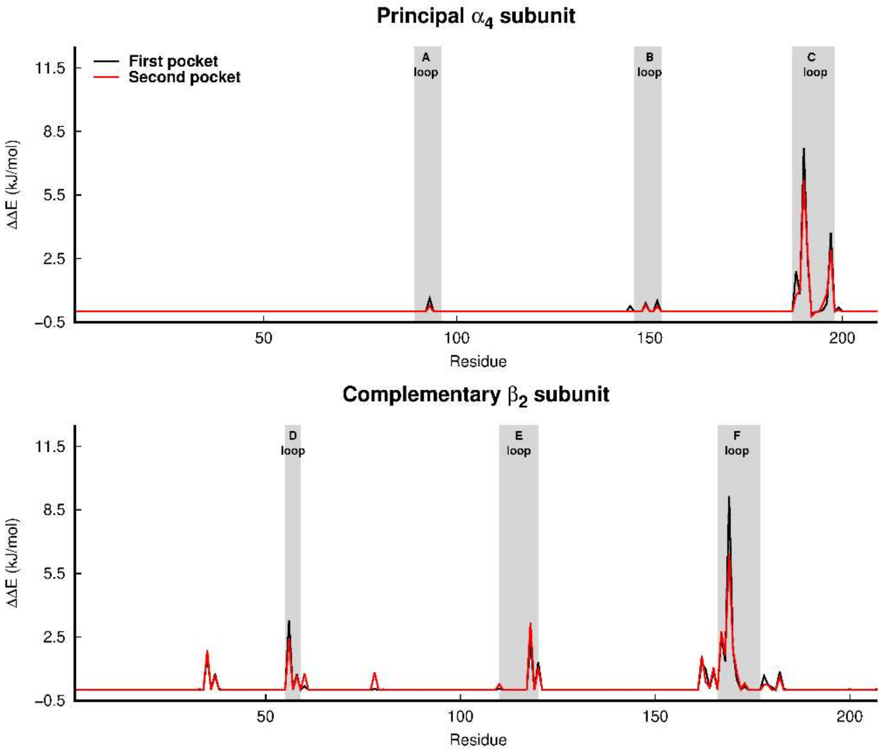

**Figure S19.** Average calculated $\Delta\Delta G_{\text{bind}}$ from the alanine-scanning mutagenesis[36] for the human α4β2 nAChR (for details see the "*In silico* alanine-scanning mutagenesis" section above). The average was determined over the three replicates. Note that the $\Delta\Delta G_{\text{bind}}$ corresponds to the difference between mutant and wild-type complexes, and as such positive $\Delta\Delta G_{\text{bind}}$ values mean that the mutation to alanine destabilizes the complex.

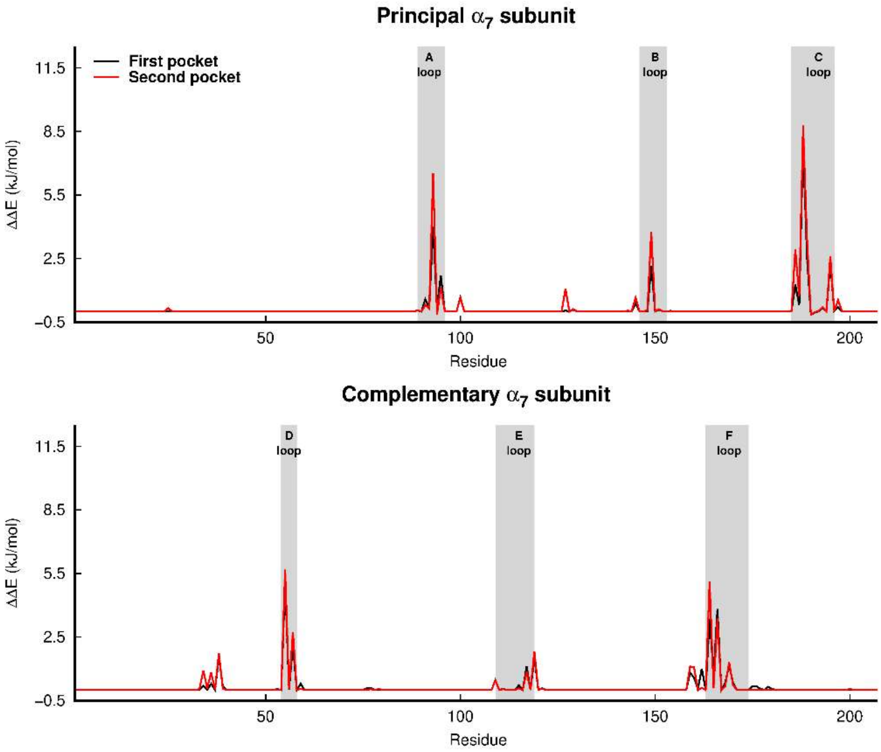

**Figure S20.** Average predicted $\Delta\Delta G_{\mathrm{bind}}$ from the alanine-scanning mutagenesis[36] for the human α7 nAChR. For more details, see the legend of **Figure S19**.

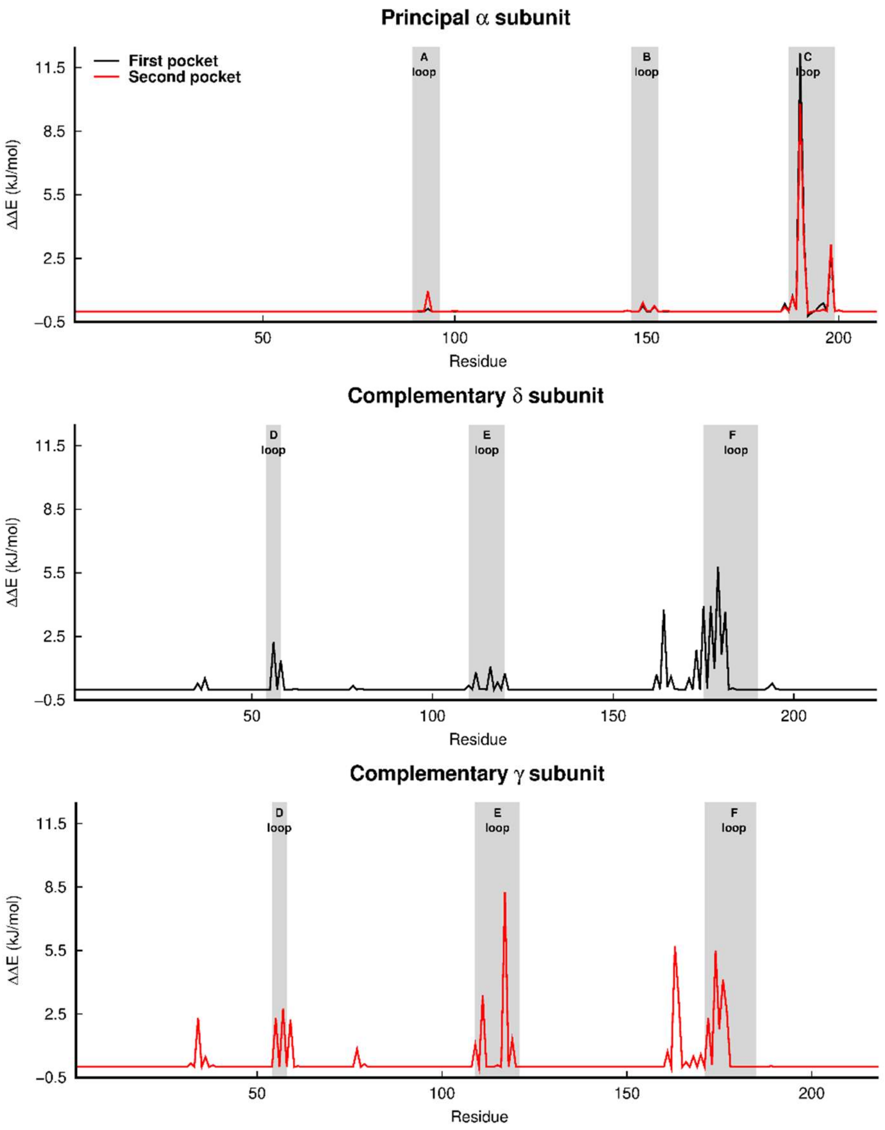

**Figure S21.** Average predicted $\Delta\Delta G_{\mathrm{bind}}$ from the alanine-scanning mutagenesis[36] for the muscle-like αβγδ nAChR from *Tetronarce californica*. For more details, see the legend of **Figure S19**.

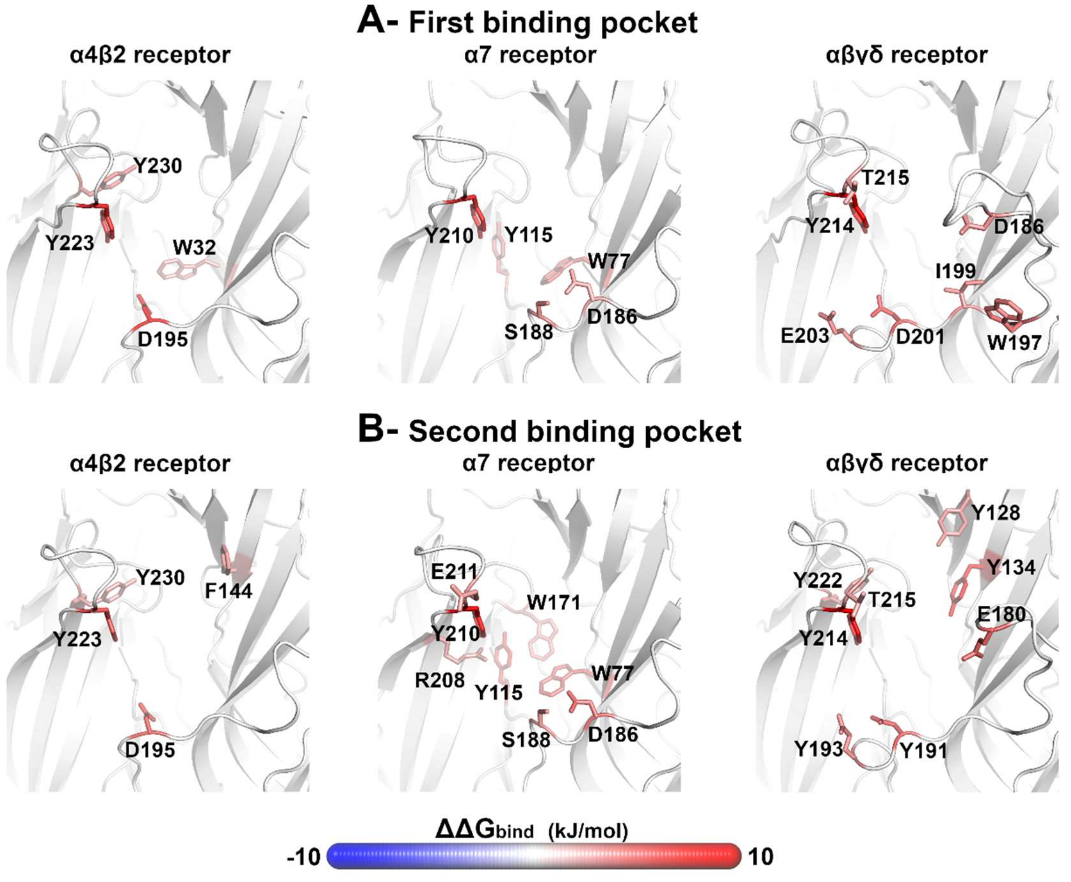

**Figure S22.** Hot spots in the binding interface of the receptors that favour binding. Note that the $\Delta\Delta G_{\text{bind}}$ corresponds to the difference between mutant and wild-type complexes. In this image, the red colour indicates a stabilizing contribution to the complex whereas blue indicates a destabilizing contribution.

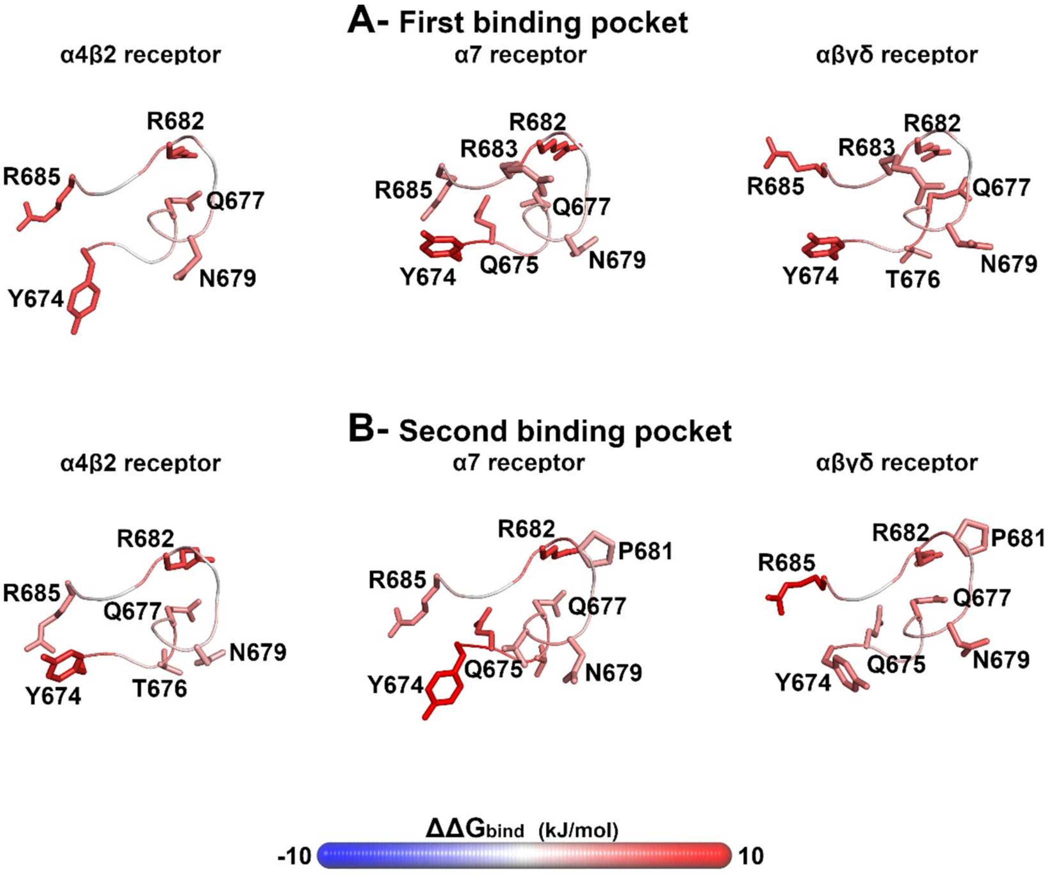

**Figure S23.** Hot spots in the S-peptide. Note that the $\Delta\Delta G_{\mathrm{bind}}$ corresponds to the difference between mutant and wild-type complexes. In this image, the red colour indicates a stabilizing contribution to the complex whereas blue indicates a destabilizing contribution.

#### Supporting tables

**Table S1:** MM-PBSA predicted relative binding energy values for the S-peptide in the human α4β2, human α7 and muscle-like αβγδ nAChR from *Tetronarce californica*. Numbers in brackets represent the standard deviations. Note that the values reported in this table do not contain the entropic contribution to the binding energy.

| | **$\Delta G_{bind}$ for the α4β2 complex** (kJ/mol) | | | |
|---|---|---|---|---|
| | **Replicate 1** | **Replicate 2** | **Replicate 3** | **Average** |
| First pocket | –308.8 (59.6) | –171.1 (51.6) | –167.9 (97.2) | –215.9 (80.4) |
| Second pocket | –274.2 (97.3) | –163.5 (92.0) | –209.5 (86.1) | –215.7 (55.7) |
| | **$\Delta G_{bind}$ for the α7 complex** (kJ/mol) | | | |
| | **Replicate 1** | **Replicate 2** | **Replicate 3** | **Average** |
| First pocket | –157.0 (78.9) | –193.2 (51.0) | –203.3 (48.0) | –184.5 (24.3) |
| Second pocket | –62.9 (72.7) | –152.8 (93.7) | –129.2 (100.6) | –114.9 (46.6) |
| | **$\Delta G_{bind}$ for the muscle-like αβγδ complex** (kJ/mol) | | | |
| | **Replicate 1** | **Replicate 2** | **Replicate 3** | **Average** |
| First pocket | –441.0 (50.2) | –420.8 (45.7) | –261.2 (73.7) | –374.3 (98.5) |
| Second pocket | –313.2 (61.0) | –464.5 (66.1) | –396.8 (81.7) | –391.5 (75.8) |

**Table S2:** Alanine-scanning predicted average $\Delta\Delta G_{bind}$ for the hot-spots (–3 kJ/mol ≥ residue contribution ≤ 3 kJ/mol) in the first binding pocket of the receptors. The average value was calculated over the three replicates. Numbers in brackets represent the standard deviations (calculated over the 303 frames per complex). Note that the $\Delta\Delta G_{bind}$ corresponds to the difference between mutant and wild-type complexes, and as such positive $\Delta\Delta G_{bind}$ values mean that the mutation to alanine destabilizes the complex.

| **First binding pocket** | | | | | |
|---|---|---|---|---|---|
| **α4β2 receptor** | | **α7 receptor** | | **Muscle-like αβγδ receptor** | |
| residue | $\Delta\Delta G_{bind}$ (kJ/mol) | residue | $\Delta\Delta G_{bind}$ (kJ/mol) | residue | $\Delta\Delta G_{bind}$ (kJ/mol) |
| β2D195 | 9.5 (3.6) | α7Y210 | 7.6 (2.2) | αY214 | 12.1 (2.6) |
| α4Y223 | 7.7 (2.0) | α7W77 | 5.1 (2.0) | δD201 | 6.1 (1.9) |
| α4Y230 | 3.7 (2.2) | α7Y115 | 3.8 (2.9) | δW197 | 4.6 (2.3) |
| β2W32 | 3.3 (1.7) | α7S188 | 3.7 (1.9) | δI199 | 4.0 (0.8) |
| | | α7D186 | 3.1 (2.1) | δD186 | 3.9 (1.7) |
| | | | | δE203 | 3.8 (2.0) |
| | | | | αT215 | 3.1 (1.5) |

**Table S3:** Alanine-scanning predicted average $\Delta\Delta G_{bind}$ values for the hot-spots (-3 kJ/mol ≥ residue contribution ≤ 3 kJ/mol) in the second binding pocket of the receptors. The average was calculated over the three replicates. Numbers in brackets represent the standard deviations (calculated over the 303 frames per complex). Note that, in this case, the $\Delta\Delta G_{bind}$ corresponds to the difference between mutant and wild-type complexes, and as such positive $\Delta\Delta G_{bind}$ values mean that the mutation to alanine destabilizes the complex.

| **Second binding pocket** | | | | | |
|---|---|---|---|---|---|
| **α4β2 receptor** | | **α7 receptor** | | **Muscle-like αβγδ receptor** | |
| residue | $\Delta\Delta G_{bind}$ (kJ/mol) | residue | $\Delta\Delta G_{bind}$ (kJ/mol) | Residue | $\Delta\Delta G_{bind}$ (kJ/mol) |
| β2D195 | 6.2 (2.9) | α7Y210 | 8.9 (1.9) | αY214 | 9.8 (2.1) |
| α4Y223 | 6.2 (2.3) | α7Y115 | 6.7 (2.4) | γY134 | 8.1 (1.7) |
| α4Y230 | 3.3 (2.6) | α7W77 | 5.6 (2.2) | γD191 | 5.7 (2.2) |
| β2F144 | 3.2 (1.6) | α7D186 | 5.3 (2.4) | γE180 | 5.5 (3.3) |
| | | α7W171 | 3.9 (1.9) | γE193 | 4.0 (2.4) |
| | | α7S188 | 3.5 (1.7) | γY128 | 3.4 (2.5) |
| | | α7E211 | 3.3 (1.4) | αY222 | 3.2 (1.4) |
| | | α7R208 | 3.0 (1.9) | αT215 | 3.0 (2.0) |
